# AAT_Rhg1_ is a tonoplast protein that alters amino acid, metabolic and defense responses and nematode resistance

**DOI:** 10.64898/2026.03.06.710000

**Authors:** Yulin Du, Aaron Lowenstein, Jorge El-Azaz Ciudad, Hiroshi A. Maeda, Andrew F. Bent

## Abstract

Soybean cyst nematode (SCN, *Heterodera glycines*) causes significant soybean yield losses. The *Rhg1* locus is a major contributor to SCN resistance and contains three genes that mediate this trait including *Rhg1-GmAAT* (*Glyma.18G022400),* which encodes putative amino acid transporter AAT_Rhg1_. The molecular function of AAT_Rhg1_ in SCN resistance is not understood. In this study, *rhg1-b* soybean lines with *Rhg1-GmAAT* silencing demonstrated that *Rhg1-GmAAT* can contribute resistance against HG 0 SCN and also against problematic HG 2.5.7 populations that partially overcome *rhg1-b*-mediated resistance. An AAT_Rhg1_ Y268L mutant complemented SCN resistance in *Rhg1-GmAAT*-silenced plants while an AAT_Rhg1_ D122A mutant did not.

Overexpression of *Rhg1-GmAAT* was not sufficient to enhance SCN resistance, suggesting that AAT_Rhg1_ requires coordinated activity with other proteins or pathways. Confocal microscopy demonstrated that AAT_Rhg1_ localizes to the tonoplast in soybean root cells. Amino acid, transcriptomic and metabolomic profiles were determined for SCN-infected root segments 3 days after SCN inoculation. In *Rhg1-GmAAT*-silenced plants relative to fully resistant (non-silenced) *rhg1-b* plants, levels of leucine, isoleucine, and tyrosine were significantly elevated. *Rhg1-GmAAT* silencing reduced SCN-responsive transcript abundances for multiple processes, significantly including genes for MAPK signaling, ethylene responses and starch and sucrose metabolism. The most common identified metabolomic changes were in amino acid derivatives, shikimate/phenylpropanoid/isoflavonoid compounds, terpenoids, and especially fatty acids. These findings can guide further investigation into the mechanisms by which AAT_Rhg1_ contributes to SCN resistance.

## Introduction

Soybean cyst nematode (SCN, *Heterodera glycines*) is responsible for substantial yield losses in major soybean production regions (Bandara *et al*., 2020; Bradley *et al*., 2021; Peng *et al*., 2021; Tylka and Marett, 2021; Júnior *et al*., 2022; Barros *et al*., 2023). During the SCN life cycle it establishes a multi-week biotrophic parasitism within soybean roots, and plant resistance may act by directly targeting the nematode or by disrupting the biotrophic interaction. Infective second-stage juveniles (J2) penetrate soybean roots and migrate to the periphery of the vascular cylinder, where they induce extensive host cell reprogramming to establish a specialized multinucleate, cytoplasm-rich feeding structure termed a syncytium (Kim *et al*., 2012; Zhang *et al*., 2017; Juvale and Baum, 2018). Fertilized females develop eggs during parasitism and subsequently form cysts that encapsulate eggs, which can remain viable in soil for several years (Hu *et al*. 2018).

Individual SCN populations are genetically diverse and can gradually overcome host resistance when soybean cultivars with similar SCN resistance are planted for multiple seasons (Acharya, Tande and Byamukama, 2016; McCarville *et al*., 2017; Gardner *et al*., 2017; Howland *et al*., 2018; Hua *et al*., 2018; Chen, 2020). SCN populations can be classified into HG Types (*Heterodera glycines* types) based on their ability to reproduce on defined soybean indicator lines that carry distinct genetic resistance sources (Niblack *et al*., 2002). For example, HG 0 populations are not capable of overcoming any of the resistance indicator lines, while HG 2 populations can partially overcome PI 88788-type resistance (Niblack *et al*., 2002). SCN populations with the same HG type can differ genetically and, even within a population, individual nematodes may have distinct mechanisms to overcome resistance.

Transcriptomic studies have explored gene expression changes in soybean roots during SCN infection for various soybean genotypes, SCN HG types, sampled tissues and time points after inoculation, with each publication providing valuable insights into the molecular interactions between soybean and SCN (Ithal *et al*., 2007a; Ithal *et al*., 2007b; Klink *et al*., 2010; Kandoth *et al*., 2011; Song *et al*., 2019; Torabi *et al*., 2023; Sultana *et al*., 2024; Wei *et al*., 2022). Transcriptomic data have been integrated with metabolomic analyses to identify altered metabolic pathways associated with susceptibility or resistance to an HG 1.2.3.5.7 SCN population (Shi *et al*., 2021).

*Rhg1* (Resistance to *Heterodera glycines* 1) is the most extensively utilized SCN resistance QTL in soybean production (Concibido *et al*., 2004; Kim *et al*., 2012; Mitchum, 2016; McCarville *et al*., 2017; Bent, 2022). *Rhg1* genes reside on a 31 to 36 kb four-gene block that exhibits DNA sequence polymorphisms and copy number variation (Cook *et al*., 2012, 2014). There are three predominant *Rhg1* haplotypes. *rhg1-a* ("Peking"-type) haplotypes often carry three tandem copies of the four-gene block*, rhg1-b (*"PI 88788"-type) haplotypes typically contain nine or ten copies, while most SCN-susceptible soybean varieties such as Williams 82 (Wm82) carry one copy of the *rhg1-c* haplotype (Cook *et al*., 2012, 2014; Lee *et al*., 2015; Liu *et al*., 2017; Patil *et al*., 2019). The basal expression level of *Rhg1* genes positively correlates with *Rhg1* copy number (Cook *et al*., 2014).

Three genes within the *Rhg1* locus have been demonstrated to contribute to SCN resistance including *Glyma.18G022400* (*Rhg1-GmAAT*), which encodes the putative amino acid transporter AAT_Rhg1_, *Glyma.18G022500 (GmSNAP18)* encoding α-SNAP_Rhg1_ (alpha-soluble NSF [N-ethylmaleimide–sensitive factor] attachment protein), and *Glyma.18G022700*, which encodes a poorly characterized protein with a wound-inducible domain (WI12_Rhg1_) (Cook *et al*., 2012; Liu *et al*., 2017; Dong & Hudson, 2022). Resistant isoforms of α-SNAP_Rhg1_ are apparently toxic to syncytia (Bayless *et al*., 2016) but further molecular mechanisms by which the three Rhg1 proteins mediate SCN resistance are not well understood (Bent, 2022).

AAT_Rhg1_, the focus of the present study, is predicted to belong to the tryptophan/tyrosine transporter protein family and contains 11 transmembrane domains (InterPro IPR013059, Pfam PF03222) (Paysan-Lafosse *et al*., 2023). AAT_Rhg1_ amino acid transport activity has not been demonstrated but indirect evidence suggests a role of AAT_Rhg1_ in modulating amino acid levels in soybean roots. Overexpression of *Rhg1-GmAAT* altered root amino acid levels, systemic glutamate transport and tolerance to exogenous glutamic acid (Guo *et al*., 2019; Du *et al*., 2025). The conserved-site mutants D122A and Y268L of AAT_Rhg1_ had opposite impacts from each other on amino acid accumulation and betalain production (Du *et al*., 2025). At the cellular level, AAT_Rhg1_ accumulated on macrovesicle-like membrane structures in SCN-penetrated cells, and the overexpression of *Rhg1-GmAAT* in soybean roots influenced membrane dynamics even in the absence of SCN infestation (Han *et al*., 2023). A recent study reported that *Rhg1-GmAAT* expression can be regulated by an AP2/ERF transcription factor GmTINY (He *et al*., 2025). Together, these findings highlight a multifaceted role of AAT_Rhg1_ in metabolic and cellular responses. Building on these observations, the present study aimed to further characterize the contribution of AAT_Rhg1_ to SCN resistance and to gain information regarding underlying mechanisms.

## Materials and Methods

More detailed descriptions of all utilized methods can be found in the Supplemental Information. A list of the oligonucleotide primers used is provided as Supplemental Table S1.

### Plasmid constructs

The *Rhg1-GmAAT* silencing construct driven by a *GmUbi* promoter is described in (Du *et al*., 2025). A synthetic artificial RNAi-escaping *Rhg1-GmAAT* coding sequence driven by a *GmUbi* promoter was created by designing synonymous codon substitutions with preference for commonly used soybean codons, synthesized at Twist Bioscience (South San Francisco, CA, USA) and deployed in binary vector pAGM4673, and also modified by site-directed mutagenesis to generate analogous *SynAATRhg1Y268L* or *SynAATRhg1D122A* constructs (Supplemental Information). Two different pAGM4673 constructs expressed wild-type AAT_Rhg1_, one native (genomic sequence spanning from 1,546 bp upstream of 5’UTR start to 440 bp downstream of 3’UTR terminus of *Rhg1-GmAAT*) and the other for constitutive overexpression (*Rhg1-GmAAT* coding sequence (exons only) controlled by a double CaMV *35S* promoter with a TMV *omega* enhancer, followed by the *nos* terminator).

For microscopy, genes encoding the fluorescent proteins mCherry, TdTomato with C-terminal KDEL endoplasmic reticulum (ER) retention signal, and mWasabi were obtained as Golden Gate level 0 plasmids. Subsequent cloning was done using the Golden Gate Assembly MoClo Tool Kit (Weber *et al*., 2011). Green fluorescent protein mWasabi was tagged to the N or C terminus of AAT_Rhg1_ in Golden Gate level 0 to level 1 reactions. The loop-tagged mWasabi AAT_Rhg1_ was created through Gibson Assembly of PCR products. *VAMP711* was cloned from Arabidopsis Biological Resource Center (ABRC) CD3-781601, and inserted into Golden Gate level 0 vector through BbsI overhangs added by PCR. The above genes were cloned into Golden Gate level 1 vectors for expression under the control of double CaMV *35S* promoter with TMV *omega* translational enhancer and *nos* terminator. The level 1 modules were assembled into the Golden Gate level 2 binary vector to generate constructs for co-expression in soybean roots.

### Transgenic plant materials

Whole transgenic *Rhg1-GmAAT*-silenced soybean plants were generated from IL3025N (*rhg1-b* SCN resistance haplotype; Haarith et al. 2026) at Wisconsin Crop Innovation Center (WCIC) as described in Du *et al*., 2025 and Supplemental Information. Transgene expression or target gene silencing were verified by RT-qPCR. For the *Rhg1-GmAAT*-silenced RNAi and corresponding EV lines, stable homozygous transgenic lines were selected, and 4th generation (T4) transgenic plants were used in SCN assays. *Rhg1-GmAAT* overexpression soybean lines were generated at WCIC. For overexpression in Williams 82 (Wm82) background, homozygous T2 plants were used in SCN assays. For overexpression in IL3025N (*rhg1-b*) and IL3849N (*rhg1-a* + *Rhg4* SCN resistance; Haarith et al. 2026), T1 plants were used in SCN assays. T-DNA copy number was tested for each T1 plant used in SCN assays to identify homozygotes, hemizygotes and non-transgenic segregants.

Composite soybean plants with transgenic roots were generated from soybean seedling hypocotyls using *Agrobacterium rhizogenes* strain K599 following the methods of Estrada-Navarrete & Alvarado-Affantranger (2007) and Fan *et al*. (2020). Detached transgenic soybean roots were produced from soybean cotyledons using the method described in Cook *et al*. (2012).

Homozygous transgenic *Arabidopsis thaliana* Col-0 lines expressing *Rhg1-GmAAT* were generated using the floral dip method (Bent, 2006). *AtAVT6C* knockout line CS864997 was ordered from ABRC.

### DNA extraction and copy number

DNA was extracted from young soybean leaf tissue using the CTAB/chloroform method. Transgene copy number was tested by qPCR, measuring the relative level of the spectinomycin resistance gene on T-DNA relative to *Glyma.18G022800*, which is known to only have one copy per haploid genome.

### SCN propagation and resistance assay

SCN populations were initially collected in the field and propagated in the laboratory on soybean plants grown in a 1:1 soil sand mixture. The HG 0 population used in experiments was originally collected in Illinois. Two HG 2.5.7 populations were sourced from Minnesota (HG 2.5.7 MN) or Missouri (HG 2.5.7 MO). To maintain or enhance virulence, these SCN populations were propagated on different soybean varieties: HG 0 on the susceptible variety Wm82 and HG 2.5.7 on *rhg1-b* variety IL3025N. Propagation pots were grown for approximately two months before harvesting. SCN cysts and eggs were extracted using the method described in Cook *et al*., 2012. Eggs were hatched in 4 mM pH 5.6 ZnCl_2_ solution.

SCN resistance assays utilized anonymized individual plants in 6cm pots containing a 1:1 soil-sand mixture, each inoculated with ∼1000 SCN eggs and incubated for 35 days, while composite plants were each inoculated with ∼750 freshly hatched SCN J2s and incubated for 30 days, prior to cyst extraction and counting. Sugar beet cyst nematode (*Heterodera schachtii*, BCN) assays on Arabidopsis were conducted as described by Butler *et al*. (2019).

### RNA extraction, RT-qPCR, and RNA-seq

RNA from flash-frozen root samples was extracted using the Direct-zol™ RNA MiniPrep Plus Kit (Zymo Research, Irvine, CA, USA; see Supplemental Information). cDNA was synthesized using FIREScript^®^ RT cDNA synthesis MIX from Solis BioDyne (Tartu, Estonia). Quantitative reverse transcription polymerase chain reaction (RT-qPCR) was performed using HOT FIREPol^®^ EvaGreen^®^ qPCR Supermix from Solis BioDyne and CFX96 real-time PCR detection system (BioRad, Hercules, CA, USA).

For RNA-seq studies (see also Supplemental Information), ten day old soybean plants grown in sand were inoculated with 1000 freshly hatched SCN (HG 0) J2s or the same volume of SCN-free hatching solution. Root segments with visible lesions caused by nematode penetration (or analogous regions from mock-inoculated plants) were flash-frozen within one minute of root collection. Four replicate samples were analyzed for each genotype-treatment. Each replicate was comprised of multiple root sections from one individual plant. RNA-seq and data analysis were performed by Novogene (Sacramento, CA, USA) using their Eukaryotic mRNA-seq service (Illumina NovaSeq platform, 150bp paired-end reads, ≥ 20 million read pairs per sample, read quality control, and gene expression quantification). The iAAT_Mock group had three replicates after one outlier was removed. Differential expression profiling primarily utilized the Novogene platform for fpkm, log2 fold change and adjusted p value (p_adj_) data, as well as KEGG and GO term functional group enrichment analysis.

### Metabolite extraction and quantification

For metabolite extraction (detailed in Supplemental Information), 12 day old soybean plants grown in sand were inoculated with SCN (HG 0) and at 3 dpi root sections were harvested, as described above. For each sample, root sections from three individual plants were pooled, flash-frozen in liquid nitrogen, and freeze-dried for 48 hours. Metabolites were extracted using a 2:1 mix of methanol and chloroform, with 50 μM of ^13^C_6_-ring-labeled-L-phenylalanine as internal standard. LC-MS analysis used a Vanquish Horizon Binary UHPLC (Thermo Scientific, Waltham, MA, USA) coupled to a Q Exactive Orbitrap mass spectrometer (Thermo Scientific, Waltham, MA, USA). Amino acids were analyzed by zwitterionic hydrophilic interaction liquid chromatography (HILIC-Z) using an InfinityLab Poroshell 120 HILIC-Z column. MS/MS settings for HILIC-Z chromatography were described in El-Azaz & Maeda, 2024. Phenylpropanoids and other semi-polar metabolites were determined by reversed-phase (RP) chromatography using an Acquity UPLC HSS T3 column, in positive ionization mode or negative ionization mode. Untargeted metabolite data processing and metabolite annotation were performed using MZmine v4.0 (Schmid *et al*., 2023) and SIRIUS v5 (Dührkop *et al*., 2019) as described in El-Azaz and Maeda, 2024. For targeted metabolite analysis, compound abundance was calculated based on the peak area of authentic standards and normalized to sample mass and internal standard recovery for each sample.

### Confocal microscopy

Confocal images were acquired using a Zeiss LSM 980 with 40x water immersion objective lens. TdTomato or mCherry signal was acquired with 561 nm laser excitation and 573 to 628 nm emission, while mWasabi signal was acquired with 488 nm laser excitation and 500 to 553 nm emission with beam splitter MBS 488/561 and laser intensity 2.0%.

### Statistical analyses

Statistical analyses (see Supplemental Information) other than RNAseq analyses were performed as described using R (version 4.4.2).

## Results

### *Rhg1-GmAAT* contributes to resistance against HG 0 and HG 2.5.7 SCN

Soybean resistance against SCN is quantitative (Bent, 2022). To test the contribution of AAT_Rhg1_ to SCN resistance in intact soybean plants and against key SCN types, we generated *Rhg1-GmAAT*-silenced transgenic soybean lines using an RNAi construct in the high-copy *rhg1-b* soybean variety IL3025N. Silencing of *Rhg1-GmAAT* was verified by RT-qPCR, with transcript levels reduced to 20-40% of the control (Supplemental Figure S1). Williams 82 (Wm82), which lacks *rhg1-b*, was used as the SCN-susceptible control to calculate the within-experiment female index (FI = 100 for Wm82), which is the most widely utilized measure of SCN resistance (Niblack *et al*., 2002). Two independent empty vector (EV) transgenic lines and two independent *Rhg1-GmAAT* RNAi (iAAT) lines were tested against three SCN populations representing two HG types. We first utilized an SCN HG 0 population originally collected in Illinois, against which *rhg1-b* remains effective. HG 0 SCN are virulent on soybeans lacking *Rhg1* resistance (e.g., Wm82). As expected, the HG 0 SCN population exhibited a substantially lower female index on *rhg1-b* EV lines than on Wm82 (Figure 1A). In the *rhg1-b* background, silencing *Rhg1-GmAAT* significantly increased the mean female index of HG 0 SCN relative to the same HG 0 SCN on EV lines, indicating compromised resistance (Figure 1A). This result is consistent with previous silencing experiments conducted on detached transgenic roots (Cook *et al*., 2012), and further supports a role for *Rhg1-GmAAT* in SCN resistance.

**Figure 1.**
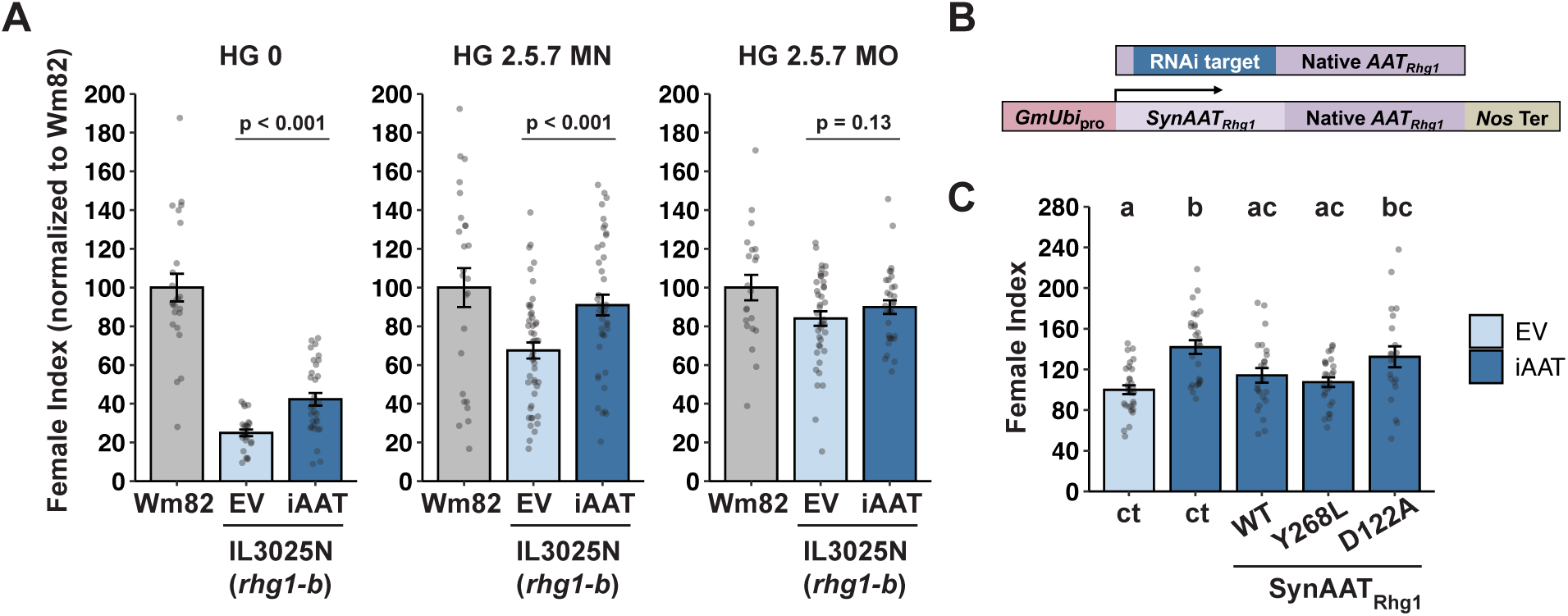
Silencing *Rhg1-GmAAT* in an *rhg1-b* background compromises SCN resistance. **(A)** SCN resistance tests with Williams 82 (Wm82) and transgenic lines, including two independent events each of IL3025N (*rhg1-b*) transformed with an empty vector (EV) or an *Rhg1-GmAAT* RNAi silencing construct (iAAT). SCN populations tested included HG 0, HG 2.5.7 from Minnesota (HG 2.5.7 MN) and Missouri (HG 2.5.7 MO). SCN resistance was measured using the Female Index (FI = 100 × [number of cysts on a tested line / number of cysts on the susceptible control Wm82]). Lower FI indicates stronger resistance. Each graph shows combined data from two independent events per transgenic genotype across two independent experiments. One-tailed *t*-tests were used to analyze the HG 2.5.7 MN and HG 2.5.7 MO experiments; Wilcoxon rank-sum test was used for HG 0 due to non-normal distribution. Data are presented as mean ± SE. Each dot represents data from an individual plant (Wm82, *n* ≥ 22; EV or iAAT, *n* ≥ 34). **(B)** Schematic of synthesized *Rhg1-GmAAT* (*SynAATRhg1*). An artificial, codon-substituted DNA fragment designed to escape the existing RNAi transgene was fused to the remaining native coding sequence to encode full-length AAT_Rhg1_. Expression was controlled by a soybean ubiquitin promoter (*GmUbi* pro) and a Nos terminator (*Nos* Ter). **(C)** Functional complementation of SCN (HG 0) resistance in composite iAAT plants expressing *SynAATRhg1*. *SynAATRhg1* (wild-type WT, Y268L, or D122A) constructs were expressed in transgenic roots generated on iAAT plants. EV and iAAT roots expressing the GFP marker alone were used as controls (ct; EV-ct IL3025N; iAAT-ct). Female Index was calculated relative to EV-ct (IL3025N). Data from two independent experiments were combined for analysis. No significant experiment effect was detected by two-way ANOVA. Bars sharing the same letter are not significantly different (Tukey’s HSD post hoc test). Data are presented as mean ± SE. Each dot represents data from an individual plant (*n* ≥ 20).

Two HG 2.5.7 SCN populations originating from Minnesota (HG 2.5.7 MN) and Missouri (HG 2.5.7 MO), each partially overcoming *rhg1-b*-mediated SCN resistance, were then tested on *Rhg1-GmAAT*-silenced lines. On EV *rhg1-b* lines, both HG 2.5.7 populations showed higher female indices than HG 0, consistent with reduced effectiveness of high-copy *rhg1-b* against HG 2.5.7 SCN (Figure 1A). Despite the elevated virulence of HG 2.5.7 MN on *rhg1-b*, silencing *Rhg1-GmAAT* resulted in significantly increased susceptibility compared with EV lines (Figure 1A), indicating that *Rhg1-GmAAT* still contributes substantially to resistance against HG 2.5.7 MN. The HG 2.5.7 MO population showed a similar trend, although not statistically significant (*t*-test *p* = 0.13; Figure 1A). The mean female index of HG 2.5.7 MO on *rhg1-b* EV lines was significantly higher than HG 2.5.7 MN (*t*-test *p* < 0.05; Figure 1A), indicating that HG 2.5.7 MO is more virulent than HG 2.5.7 MN on a full *rhg1-b* background, even though both are classified as HG 2.5.7 populations.

A transgenic complementation test further validated the role of *Rhg1-GmAAT* in SCN resistance. We generated an artificial codon-substituted, RNAi-escaping *Rhg1-GmAAT* (*SynAATRhg1WT*; Figure 1B). *SynAATRhg1WT* encodes a functional protein, as it enhanced betalain accumulation when co-overexpressed with *RUBY*, consistent with a prior study (He *et al*., 2020; Du *et al*. 2025) (Supplemental Figure S2). For the complementation test, *SynAATRhg1WT* was overexpressed via a soybean ubiquitin promoter in transgenic roots of composite soybean plants derived from the *Rhg1-GmAAT*-silenced (iAAT) lines described above. When inoculated with SCN HG 0, overexpression of *SynAATRhg1WT* in the iAAT background restored resistance to levels equivalent to *rhg1-b* EV plants (EV-ct; Figure 1C), indicating that the compromised resistance in the RNAi lines was successfully complemented.

### Functional consequences of AATRhg1 D122A and Y268L mutations in SCN resistance

In a recent study we showed that soybean AAT_Rhg1_ Y268L and D122A mutations, at residues that are highly conserved across plant amino acid transporters, differentially affect AAT_Rhg1_ function in regulating amino acid homeostasis and *RUBY*-mediated betalain synthesis (Du *et al*., 2025). The D122A mutation apparently blocked AAT_Rhg1_ function, while the AAT_Rhg1_Y268L mutant was postulated to have increased activity as it induced a stronger betalain accumulation than wild-type AAT_Rhg1_ (Du *et al*., 2025). To examine whether these two mutants also affect AAT_Rhg1_-mediated SCN resistance, *SynAATRhg1Y268L* or *SynAATRhg1D122A* variants were overexpressed in *Rhg1-GmAAT* RNAi lines and tested against SCN HG 0. In transgenic roots, synthesized and/or native *Rhg1-GmAAT* transcript levels were measured by RT-qPCR (Supplemental Figure S3). The Y268L mutant restored the compromised SCN resistance phenotype in the iAAT background, but did not confer greater resistance than *SynAATRhg1WT* (Figure 1C). As predicted for a loss-of-function mutation, the D122A mutant did not differ significantly from iAAT control (iAAT-ct) plants, indicating that D122A failed to complement SCN resistance (Figure 1C).

### Overexpressing *Rhg1-GmAAT* alone does not enhance SCN resistance

We previously reported that *rhg1-a*, *rhg1-b* or *rhg1-c* encode identical AAT_Rhg1_ proteins, and that increased *Rhg1* copy number correlates with elevated *Rhg1-GmAAT* transcript levels and enhanced SCN resistance (Cook *et al*., 2014). In resistant soybean varieties, AAT_Rhg1_ protein abundance is further induced upon SCN infestation and accumulates along the SCN penetration path (Han *et al*., 2023). In the present study we transgenically expressed *Rhg1-GmAAT* in various soybean genetic backgrounds to test whether constitutively elevated *Rhg1-GmAAT* expression enhances SCN resistance. A *35S promoter:Rhg1-GmAAT CDS:Nos terminator* construct was transformed into SCN-susceptible single-copy *rhg1-c* Wm82 and SCN-resistant low-copy *rhg1-a* variety IL3849N. A *native promoter:native Rhg1-GmAAT (UTR and introns included):native terminator* construct was transformed into Wm82 and SCN-resistant high-copy *rhg1-b* variety IL3025N. *Rhg1-GmAAT* transcript levels in independent transgenic lines were quantified by RT-qPCR (Supplemental Figure S4). Transgene presence significantly elevated *Rhg1-GmAAT* transcript levels in most lines, but with the native construct, two lines (Wm82 Line 2 and *rhg1-b* Line 2) showed only marginal increases and one line (Wm82 Line 1) showed no significant change (Supplemental Figure S4). Transgenic lines were then tested against SCN populations chosen for substantial virulence on the corresponding *rhg1* genetic backgrounds. Elevated *Rhg1-GmAAT* transcript levels did not result in significantly enhanced SCN-resistance relative to non-transformed plants or non-transgenic segregant controls (NT_seg_) in any tested genotype (Figure 2). These results suggest that increasing *Rhg1-GmAAT* transcript abundance to the extent we generated alone is insufficient to further enhance SCN resistance.

**Figure 2.**
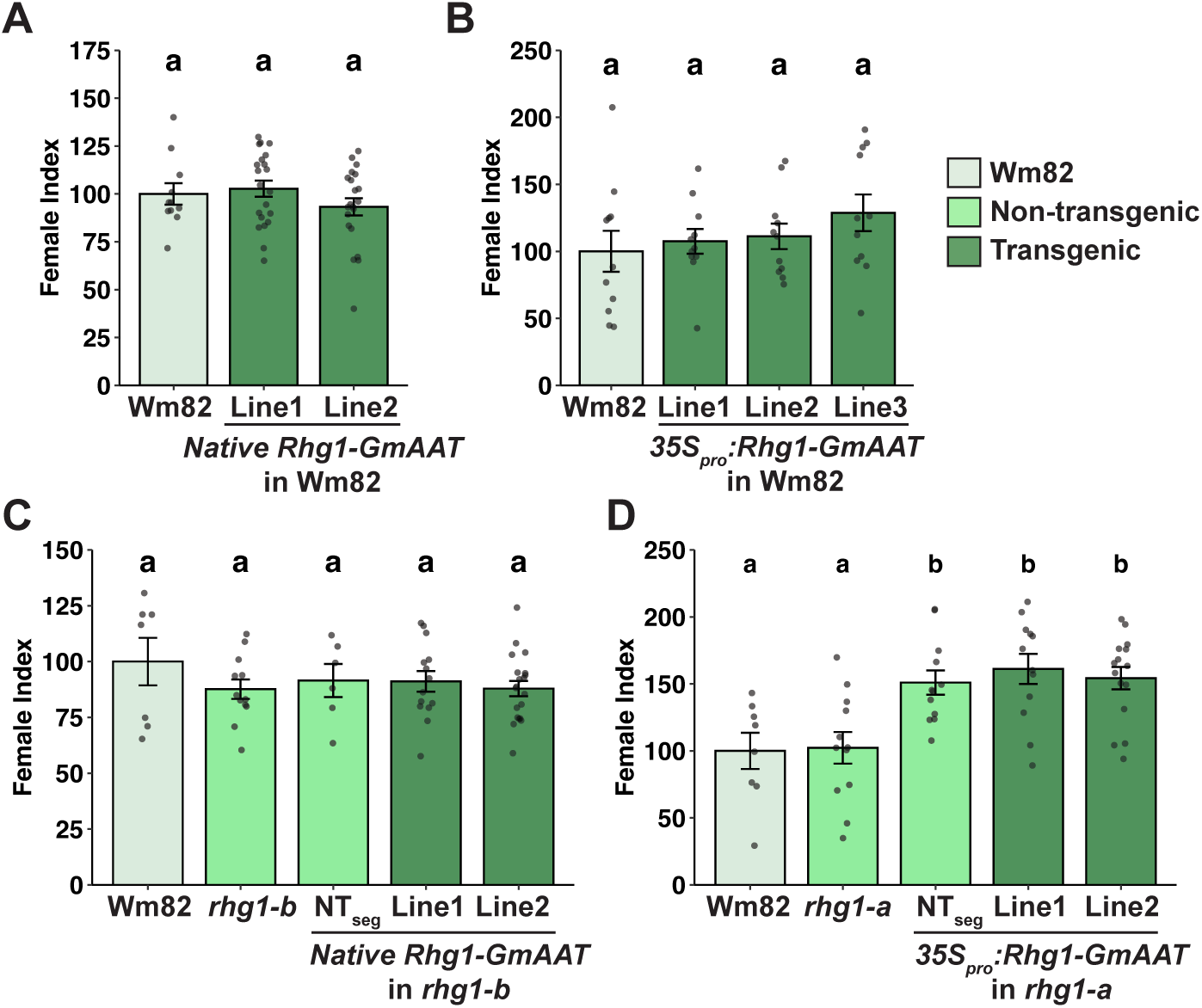
Overexpression of *Rhg1-GmAAT* does not enhance SCN resistance. Soybean lines transgenically expressing *Rhg1-GmAAT* were tested against compatible SCN populations. *Rhg1-GmAAT* transcript levels for each line are shown in Supplemental Figure S4. Unless otherwise noted, transgenic lines exhibited significantly elevated *Rhg1-GmAAT* transcript levels relative to non-transgenic controls. **(A)** SCN HG 0 resistance test on Wm82 transformed with a native *Rhg1-GmAAT* construct (native promoter, UTRs, exons, introns and terminator). Line 2 showed an 11-fold but marginally significant increase (*p* = 0.1) in transcript level. **(B)** SCN HG 0 resistance test on Wm82 transformed with an overexpression construct (*35S promoter:Rhg1-GmAAT CDS*). **(C)** SCN HG 2.5.7 resistance test on IL3025N (*rhg1-b*) soybean transformed with the native construct. Controls include non-transgenic *rhg1-b* and segregated non-transgenic progeny from transformed plants (NT_seg_). Line 1 showed no increase in *Rhg1-GmAAT* expression, and Line 2 exhibited a 9-fold but marginally significant increase (*p* = 0.07). **(D)** SCN HG 1.3.6.7 resistance test on IL3849N (*rhg1-a*) soybean transformed with the *35S* overexpression construct, with non-transgenic and NT_seg_ controls. Female index was calculated relative to Wm82. Data were analyzed using ANOVA with post hoc Tukey’s HSD. Bars sharing the same letter within a graph are not significantly different. Data are presented as mean ± SE. Each dot represents data from an individual plant.

### Overexpression of *Rhg1-GmAAT* or loss of *AtAVT6C* in Arabidopsis does not alter BCN resistance

A previous study showed that co-overexpression of the three soybean *Rhg1* genes in Arabidopsis enhanced resistance against BCN (Butler *et al*., 2019). To determine whether overexpression of *Rhg1-GmAAT* alone could confer BCN resistance in Arabidopsis, we transformed Arabidopsis Col-0 plants with the two *Rhg1-GmAAT* expression constructs described above. Stable homozygous transgenic lines were selected and *Rhg1-GmAAT* transcript levels were tested (Supplemental Figure S5A). These transgenic Arabidopsis lines did not show differences in BCN susceptibility compared with empty vector control lines (Supplemental Figure S5B).

*AtAVT6C* is the ortholog of *Rhg1-GmAAT* in *Arabidopsis thaliana*. A Col-0 *AtAVT6C* Arabidopsis knockout line was obtained and validated by PCR-based genotyping (Supplemental Figure S6A). Subsequent BCN infection assays showed no significant differences in susceptibility between the knockout mutant and wild-type Col-0 accession (Supplemental Figure S6B).

### AATRhg1 protein colocalizes with tonoplast marker VAMP711 in living soybean root cells

AAT_Rhg1_ protein was predicted to have a vacuolar membrane (tonoplast) or cell membrane localization by DeepLoc-2.1 (Ødum *et al*., 2024), with its N terminus facing the cytosol according to DeepTMHMM prediction (Hallgren *et al*., 2022). To experimentally validate the tonoplast location of AAT_Rhg1_ suggested in our preliminary microscopy experiments, mWasabi-tagged AAT_Rhg1_ was coexpressed in soybean root cells with a frequently used tonoplast marker, mCherry-tagged VESICLE-ASSOCIATED MEMBRANE PROTEIN 711 (VAMP711) (Geldner *et al*., 2009). The tonoplast localization of mCherry-VAMP711 was verified in soybean roots (Supplemental Figure S7). In root regions with nearly fully elongated cells, the tonoplast was clearly distinguished from the plasma membrane by the surrounding soluble cytosolic mWasabi (Figure 3A). AAT_Rhg1_ was tagged with mWasabi either on the loop between the transmembrane helix 6 (TM6) and TM7 (AAT_Rhg1_ loop mWasabi) or on its N terminus (AAT_Rhg1_ N-mWasabi). In soybean roots, AAT_Rhg1_ loop mWasabi primarily colocalized with mCherry-VAMP711 to the tonoplast, indicating a vacuole membrane localization (Figure 3A). AAT_Rhg1_ N-mWasabi also showed partial colocalization with mCherry-VAMP711 but substantially colocalized with TdTomato fused to an endoplasmic reticulum retention signal (TdTomato-ER), suggesting that N-terminal tagging of AAT_Rhg1_ can affect its subcellular trafficking (Figure 3B). Non-transgenic soybean roots and roots expressing only mCherry or mWasabi were imaged as controls for autofluorescence and channel bleed-through (Supplemental Figure S7).

**Figure 3.**
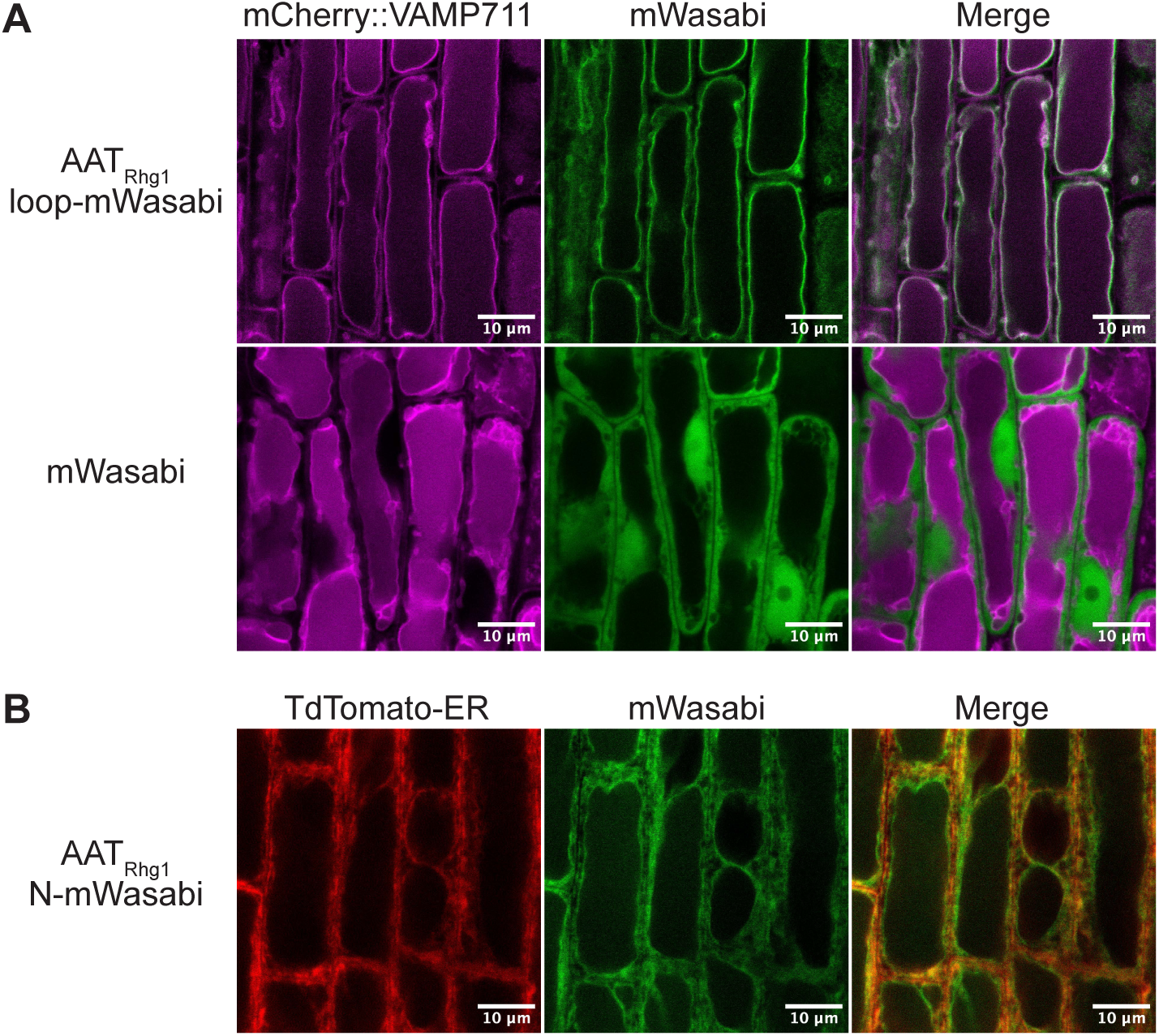
GFP-tagged AATRhg1 protein localizes to the tonoplast in soybean root cells. **(A)** Representative confocal microscopy images of transgenic soybean roots co-expressing the tonoplast marker mCherry::VAMP711 (magenta) and either AAT_Rhg1_ with mWasabi inserted in the loop between transmembrane helices TM6 and TM7 (loop-mWasabi, green) or a soluble mWasabi control (green). Images shown are from the root zone containing nearly fully elongated root cells. White pixels in the merged images indicate colocalization. Scale bars = 10 µm. **(B)** Representative confocal microscopy images of transgenic soybean root co-expressing TdTomato with an endoplasmic reticulum retention signal (TdTomato-ER, red) and N-terminally mWasabi-tagged AAT_Rhg1_ (AAT_Rhg1_ N-mWasabi, green). Colocalization appears yellow in the merged channel. Scale bars = 10 µm.

### Silencing *Rhg1-GmAAT* affects amino acid homeostasis

Overexpression of *Rhg1-GmAAT* has been shown to affect amino acid homeostasis in soybean roots (Guo *et al*., 2019; Du *et al*., 2025). To examine the impact of silencing *Rhg1-GmAAT* on amino acid levels, the *rhg1-b* EV or iAAT plants described above were mock-inoculated or inoculated with HG 0 SCN, and 18 amino acids were quantified from infected root sections at 3 dpi using HILIC-Z liquid chromatography-mass spectrometry (LC-MS) (Figure 4, Supplemental Figure S8). In the absence of SCN, silencing *Rhg1-GmAAT* resulted in significantly higher levels of leucine, isoleucine and tyrosine (Figure 4). In the presence of SCN at 3 dpi, isoleucine and tyrosine levels were significantly higher in iAAT roots than in EV roots (Figure 4; p = 0.66 for leucine). Phenylalanine and methionine showed a similar but non-significant trend (Supplemental Figure S8). Other quantified amino acids showed no significant differences between EV and iAAT plants (Supplemental Figure S8). Aspartic acid and glutamic acid levels increased in response to SCN infection in both genotypes, with no observed impact from silencing *Rhg1-GmAAT* (Supplemental Figure S8; similar trend for Asn and Pro not statistically significant). These findings further indicate that AAT_Rhg1_ protein abundance affects certain, but not all, amino acid levels in soybean roots.

**Figure 4.**
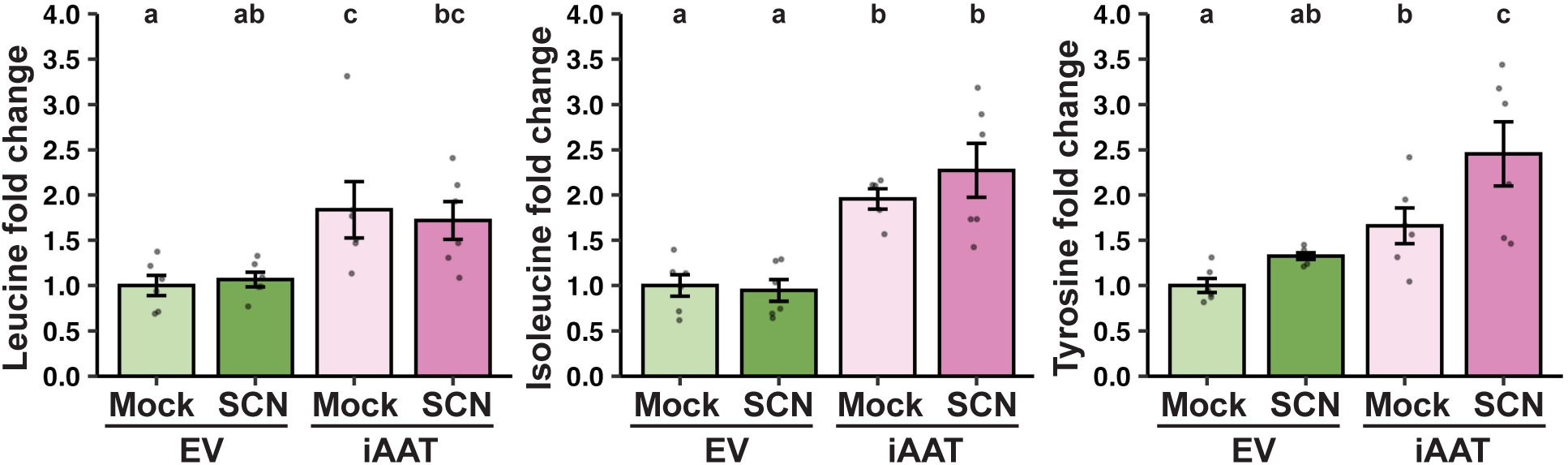
Silencing *Rhg1-GmAAT* in an *rhg1-b* background alters specific amino acid levels. Quantification of amino acid levels in roots of IL3025N (*rhg1-b*) empty vector (EV) or *Rhg1-GmAAT* RNAi-silenced (iAAT) transgenic soybean lines, either mock-treated or inoculated with SCN (HG 0). Samples were collected 3 days post-inoculation (dpi). Amino acid levels were normalized as fold change relative to EV-Mock samples within the same experiment. Data from two independent experiments were combined for analysis. Data for each amino acid were analyzed using a linear model including group and replicate as fixed effects. Bars sharing the same letter within a graph are not significantly different. Data are presented as mean ± SE (*n* = 6).

### Transcriptomic analysis reveals altered responses to SCN when *Rhg1-GmAAT* is silenced

To better understand the role of *Rhg1-GmAAT* in SCN resistance, transcriptomic analyses were performed on the *rhg1-b* EV and iAAT lines described above. Plants were either inoculated with an HG 0 SCN population or mock-inoculated. Root tissues were collected at 3 days post-inoculation (dpi) from ∼0.5 cm root sections exhibiting visible SCN infection site lesions in inoculated plants, or from corresponding root regions in mock-inoculated controls. Three dpi was chosen as a time when syncytia are established and expanding, 2-3 days prior to the visible syncytial collapse caused by the *rhg1-b* resistance trait (Kim *et al*., 2010, 2012; Mitchum, 2016). Differentially expressed genes (DEGs) were identified using an adjusted *p*-value threshold of *p*_adj_ ≤ 0.05, with *p*-values adjusted using the Benjamani-Hochberg method to control the false discovery rate. Because SCN directly impact a minority of the cells within infected root samples, no minimum transcript abundance fold-change cutoff was imposed to reduce false-negative elimination of valid DEGs whose transcript changes may be diluted by surrounding less-affected cells.

DEGs first were identified between iAAT-Mock and EV-Mock samples to investigate transcriptional changes caused by silencing *Rhg1-GmAAT* in the absence of SCN. Silencing *Rhg1-GmAAT* resulted in only 104 significantly upregulated and 78 significantly downregulated genes, suggesting minimal pleiotropic effects from reduced *Rhg1-GmAAT* expression (Supplemental Figure S9A, Supplemental Table S2). None of the protein-coding DEGs corresponded to known core enzymes directly involved in the biosynthesis or catabolism of leucine, isoleucine or tyrosine. KEGG analysis revealed limited pathway enrichment among these DEGs, with only one group enriched to a statistically significant extent: a number of MAPK signaling pathway genes were significantly upregulated in iAAT-Mock relative to EV-Mock roots, all of which were associated with ethylene signaling (Supplemental Figure S9B, Supplemental Table S3 tab for Fig. S9C).

In contrast to the mock-only comparison above, more than five thousand DEGs were observed upon SCN infection within each genotype (SCN vs. Mock comparisons; Supplemental Figure S10A and S10B, Supplemental Table S2). KEGG enrichment analyses indicated broad transcriptional responses to SCN infection (Supplemental Figure S10C to S10F, Supplemental Table S3). For infected root areas of SCN-resistant *rhg1-b* EV plants, observed transcriptional SCN responses (Supplemental Figure S10) carried multiple similarities to previously reported SCN-responsive gene expression profiles (Kandoth *et al*., 2011; Song *et al*., 2019; Wei *et al*., 2022; Zhang *et al*., 2022; Torabi *et al*., 2023; Sultana e*t al.*, 2024). KEGG pathways significantly enriched among SCN-upregulated genes in *rhg1-b* EV roots but not in iAAT roots included MAPK signaling, protein processing in ER, cyanoamino acid metabolism, starch and sucrose metabolism, and other groups (asterisks in Supplemental Figure S10C).

To more stringently examine SCN-responsive transcriptional differences between *rhg1-b* EV and iAAT plants, we refined the above SCN-responsive DEG lists by separating out the 2,160 upregulated and 1,007 downregulated genes shared by both plant genotypes (Figure 5A-D, Supplemental Table S2). Those shared DEGs were enriched in KEGG pathways related to defense and metabolism, as well as circadian rhythm, DNA replication and protein processing (Figure 5C and 5D, Supplemental Table S3), suggesting that *Rhg1-GmAAT* is less central in modulating those SCN responses. Excluding the shared DEGs allowed identification of SCN-responsive genes whose expression is robustly affected by *Rhg1-GmAAT* silencing, which are listed in Supplemental Table S2 on tabs whose names start with the word “Unique”. Their significantly overrepresented KEGG groups are presented in Figure 5E-H. It remained of particular interest to identify genes upregulated in response to SCN in *rhg1-b* EV roots but not in iAAT roots. This set of genes was again enriched for MAPK signaling pathway, starch and sucrose metabolism, and a pair of overlapping KEGG gene sets related to secondary metabolite biosynthesis and cyanoamino acid metabolism (Figure 5E; individual genes listed in Supplemental Table 3 tab for Fig. 5E). Intriguingly, no KEGG groups were significantly over-represented among EV-specific downregulated genes (Figure 5F). In the more SCN-susceptible iAAT roots, genes uniquely upregulated during SCN infection were associated with motor proteins, proteasome, and DNA replication (Figure 5G), whereas genes uniquely downregulated in iAAT roots were significantly enriched in metabolic and stress-response pathways: MAPK signaling, branched-chain amino acid (Val/Leu/Ile) degradation, carotenoid biosynthesis, and autophagy (Figure 5H).

**Figure 5.**
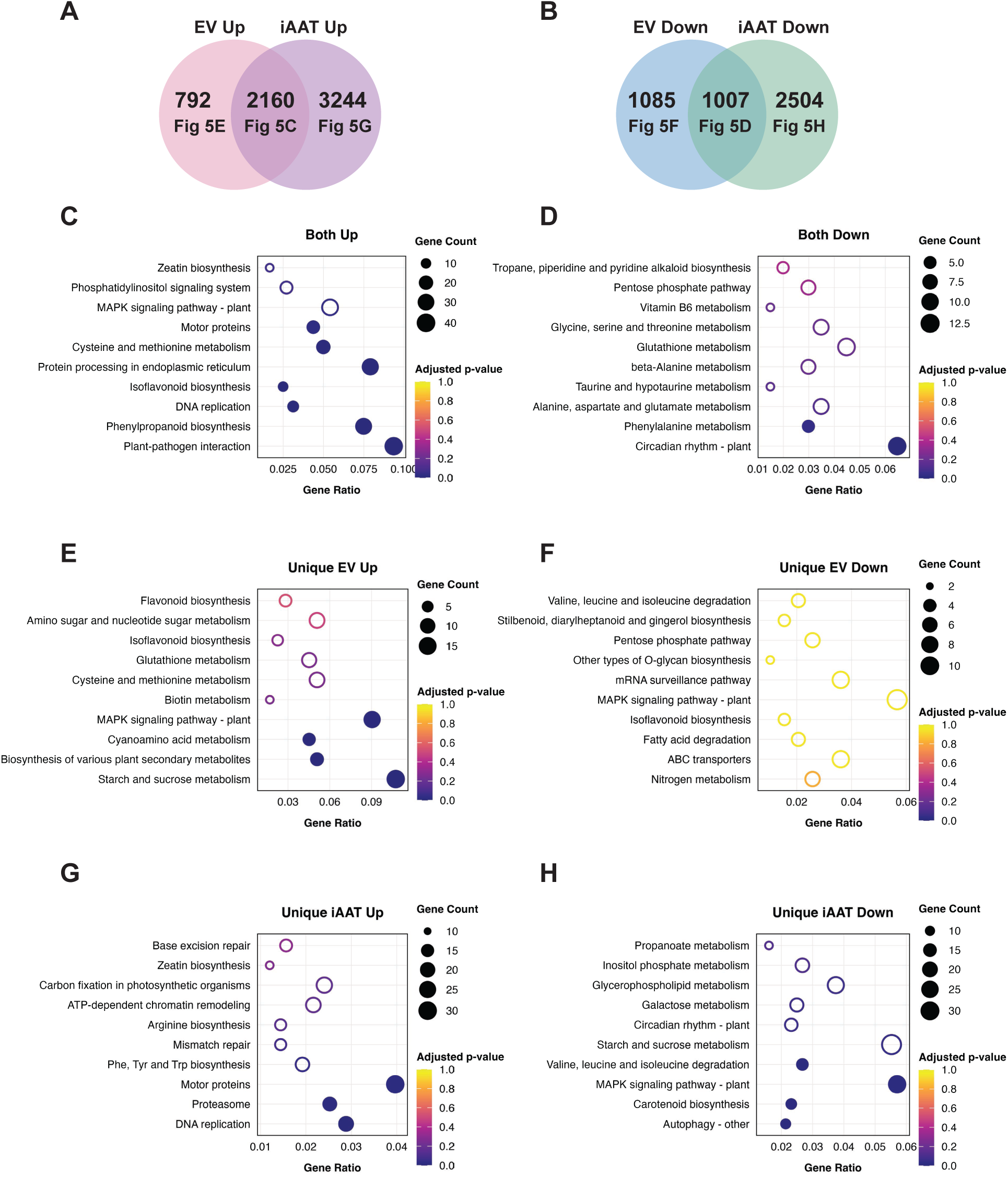
Shared and genotype-specific transcriptional responses to SCN infection in EV and iAAT soybean lines. **(A, B)** Venn diagrams showing overlaps of SCN-responsive differentially expressed genes (DEGs) between EV and iAAT lines. The numbers of upregulated genes are shown in **(A)**, and downregulated genes in **(B)**. **(C–H)** KEGG pathway enrichment dot plots for DEGs shared between or unique to genotypes. DEGs shared by both genotypes are shown in **(C)** for upregulated genes and **(D)** for downregulated genes. EV-specific DEGs are shown in **(E)** for upregulated and **(F)** for downregulated genes. iAAT-specific DEGs are shown in **(G)** for upregulated and **(H)** for downregulated genes. Dot size represents the number of DEGs associated with each pathway, and color indicates enrichment significance. Filled circles illustrate significantly enriched pathways (adjusted *p*-value < 0.05); empty circles indicate non-significant enrichment.

Given that MAPK signaling pathway genes (KEGG ID gmx04016) were significantly overrepresented as being upregulated in response to SCN infection uniquely in SCN-resistant *rhg1-b* EV roots, and downregulated uniquely in iAAT roots, we further examined this finding and discovered a high presence of ethylene-associated genes. Nine of the 16 genes on the MAPK signaling pathway KEGG list for the “Unique EV Up” set of DEGs are annotated for ethylene synthesis, perception or responses (Figure 6, Supplemental Table 3 tab for Fig. 5E). An additional 11 ethylene-associated genes were also in that “Unique EV Up” DEG group but are not annotated as part of the KEGG MAPK signaling pathway (Figure 6). Intriguingly, in the “Unique iAAT Down” set, 16 of 32 genes DEGs in the KEGG MAPK signaling pathway (Fig. 5H) were also ethylene-associated genes, as were an additional 24 genes in the “Unique iAAT Down” DEG set (Supplemental Figure S11; Supplemental Table S3 tab for Fig. 5H). AAT_Rhg1_ apparently makes a significant contribution to *rhg1-b* ethylene-mediated responses to SCN infection.

While the above KEGG analyses identify broad pathway trends, it should be noted that multiple individual genes with potentially high biological relevance were not highlighted. The complete list of DEGs is provided in Supplemental Table S2.

### Untargeted metabolomic profiling indicates broad metabolic changes in *Rhg1-GmAAT*-silenced soybean roots

To investigate metabolomic changes associated with *Rhg1-GmAAT* silencing, we performed untargeted metabolomic analysis on root samples from EV and iAAT plants, either mock-inoculated or inoculated with SCN at 3 dpi. Principal component analysis (PCA) showed that silencing of *Rhg1-GmAAT* substantially affected the levels of detectable metabolites, with a stronger overall effect on metabolomic profiles than SCN infection itself (Figure 7A).

**Figure 6.**
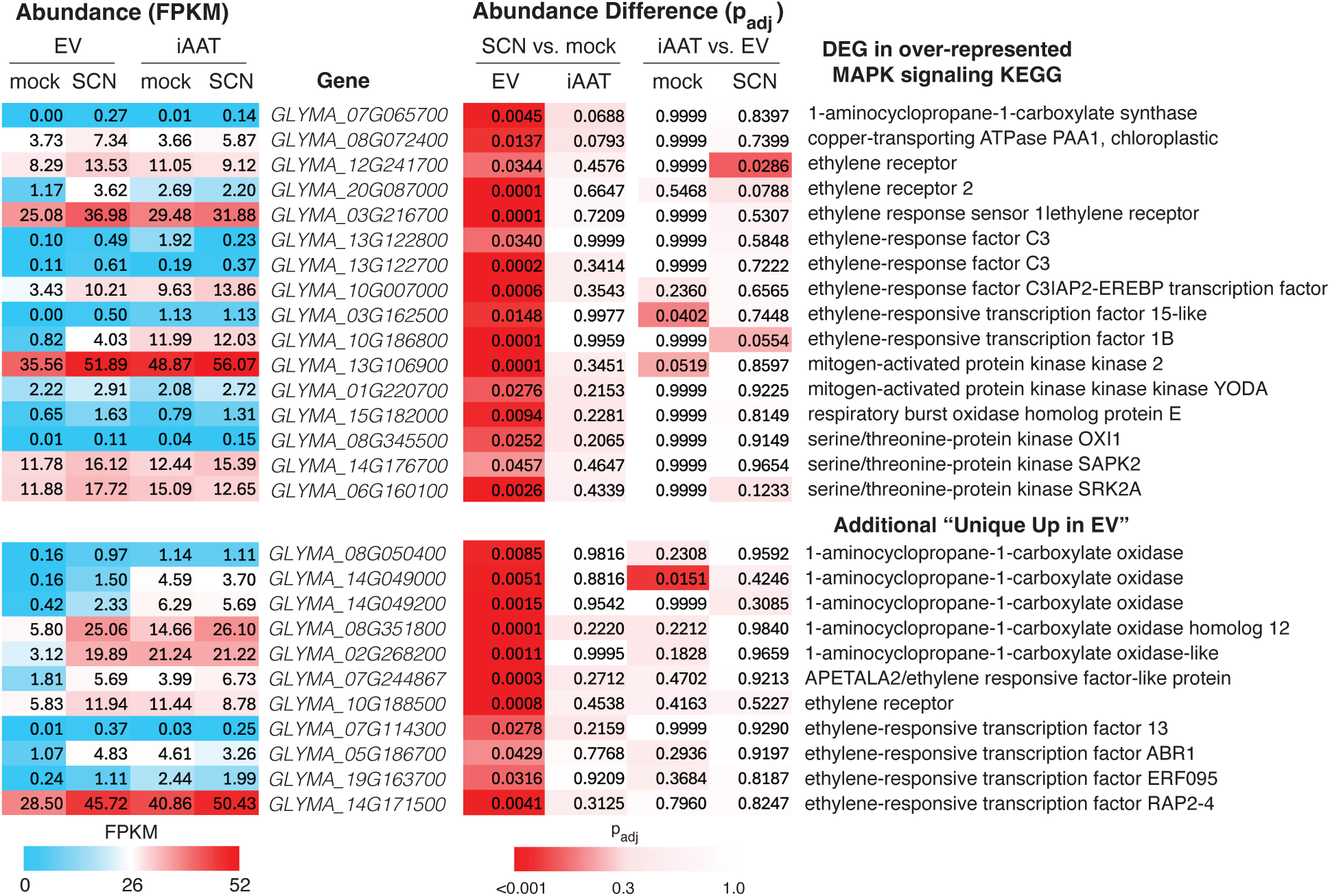
Silencing of *Rhg1-GmAAT* significantly alters abundance of transcripts associated with MAPK signaling and ethylene responses. Top: Transcripts significantly elevated at SCN infection sites in SCN-resistant *rhg1-b* (EV) roots but not in *Rhg1-GmAAT*-silenced roots (iAAT), for over-represented KEGG group “MAPK Signaling Pathways – Plants”. Bottom: Additional genes uniquely up in EV but not iAAT roots in response to SCN, that carry ethylene-associated annotation but are not placed by KEGG into the above group. See also Supplemental Figure S11 showing 40 ethylene-associated genes and other MAPK signaling genes uniquely down in iAAT but not EV.

**Figure 7.**
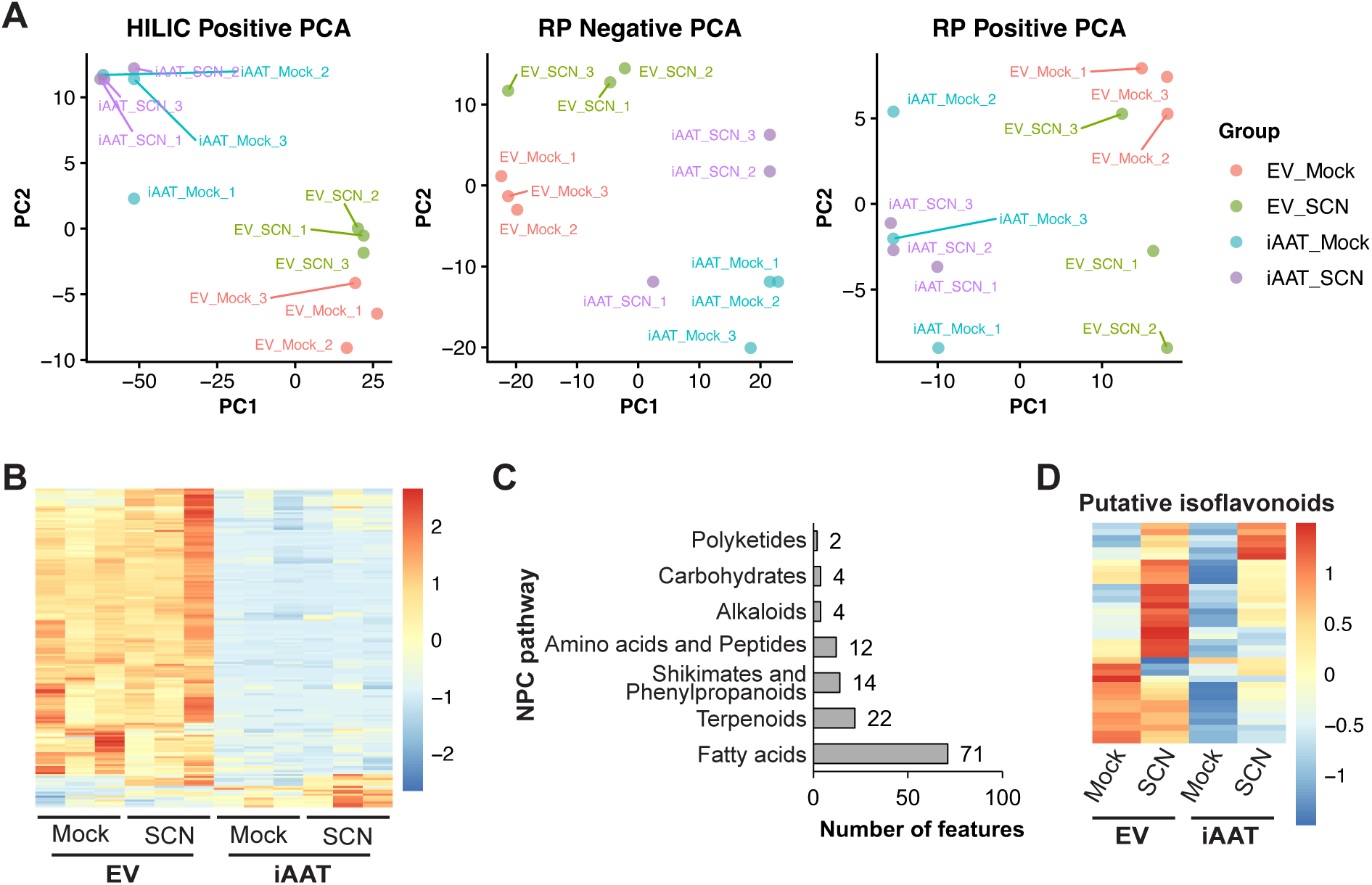
Silencing *Rhg1-GmAAT* in an *rhg1-b* background alters metabolomic profiles. **(A)** Principal component analyses (PCA) of untargeted metabolomic profiles obtained using hydrophilic interaction chromatography (HILIC, positive mode) and reversed-phase chromatography (RP, positive and negative modes). Metabolites were quantified from infection zones from roots of *rhg1-b* EV or iAAT transgenic soybean lines, either mock-inoculated or inoculated with SCN (HG 0) at 3 dpi. **(B)** Heatmap showing mass features that were significantly altered (fold-change > 2 or < 0.5; *t*-test *p* < 0.05) in at least one pairwise comparison between EV-mock vs. iAAT-mock or EV-SCN vs. iAAT-SCN. Colors represent relative metabolite abundance detected with RP-negative mode. **(C)** NPC pathway classification of the differentially abundant metabolites in (B) with available NPC annotations. Bars indicate the number of metabolites assigned to each NPC biosynthetic pathway. **(D)** Heatmap showing the relative abundances of predicted isoflavonoids detected in RP-negative mode.

To further characterize these metabolic shifts, we analyzed LC-MS data using MZmine (Schmid *et al*., 2023), followed by molecular structure prediction in SIRIUS (Dührkop *et al*., 2019). Because reversed-phase (RP) negative mode ionization yielded overall higher compound prediction confidence, subsequent analyses focused on this dataset. In total, 612 mass features were detected, of which 412 were assigned predicted molecular structures (Supplemental Table S4). Among all detected features, 162 (26%) showed significant abundance changes (fold-change > 2 or < 0.5; *t*-test *p* < 0.05) in at least one pairwise comparison between EV-mock vs. iAAT-mock or EV-SCN vs. iAAT-SCN (Figure 7B). To gain insight into the metabolic categories affected by *Rhg1-GmAAT* silencing, annotated compounds were classified using the Natural Products Classification (NPC) system (Kim *et al*., 2021). Among the significantly altered features, fatty acids represented the largest category, followed by terpenoids, shikimates and phenylpropanoids, and amino acid derived metabolites (Figure 7C, Supplemental Figure S12).

Isoflavonoids have been implicated previously in soybean resistance to SCN (Jiang *et al*., 2025), and genistein and daidzein are recognized as soybean phytoalexins due to their antimicrobial activities (Dakora and Phillips, 1996). Analysis of predicted isoflavonoid-related features from the RP negative mode dataset showed an overall trend towards reduced abundance in iAAT roots compared with EV roots, in both mock-treated and SCN-infected tissues, with particularly low isoflavonoid levels in mock-treated iAAT roots (Figure 7D; Supplemental Table S4).

## Discussion

Our results show that *Rhg1-GmAAT* can contribute to resistance against not only HG 0, but also a more virulent HG 2.5.7 SCN population that partially overcomes the resistance conferred by *rhg1-b*. The HG 2.5.7 MN population may have evolved mechanisms to escape the quantitative resistance conferred via the other two *Rhg1*-encoded proteins but not AAT_Rhg1_. Alternatively, it may partially overcome the resistance conferred by AAT_Rhg1_, but in either case, AAT_Rhg1_ still provides quantitative resistance.

Because of genetic heterogeneity within SCN populations, it is also possible that only some individuals are capable of overcoming AAT_Rhg1_. The observed differences between HG 2.5.7 MN and HG 2.5.7 MO populations further suggest that SCN populations classified under the same HG type may have evolved through distinct paths, leading to variations in their ability to reproduce on a resistant soybean line.

Overexpression of *Rhg1-GmAAT* alone, regardless of SCN resistance background, was insufficient to enhance resistance as measured in the present study. A previous, more limited set of experiments also failed to detect increased resistance from *Rhg1-GmAAT* overexpression (Liu *et al*., 2017). In contrast, elevated expression of α-SNAP_Rhg1_LC (*GmSNAP18*) has been shown to quantitatively enhance resistance against SCN (Liu *et al*., 2017; Haarith *et al*., 2025). Co-overexpression of the three *Rhg1* genes in soybean has been shown to promote resistance against SCN in transgenic Wm82 roots (Cook *et al*., 2012). Beyond soybean, transformation of Arabidopsis and potato plants with constructs that direct overexpression of the three *Rhg1* genes also elevated resistance, in that case against BCN and potato cyst nematodes (*Globodera rostochiensis* and *Globodera pallida*) (Butler *et al*., 2019).

Apparently, AAT_Rhg1_ itself is necessary but not sufficient for full resistance. These findings support the hypothesis that the three *Rhg1* genes function synergistically, with all three required to trigger full SCN resistance. It is possible that AAT_Rhg1_ contributes to a broader physiological or metabolic context that facilitates resistance responses, such as maintaining amino acid or phytohormone homeostasis, priming phytoalexin biosynthesis or activation of defense signaling cascades. These pathways may not be fully engaged without coordinated tissue- and infection-appropriate expression of AAT_Rhg1_ along with the other two *Rhg1* proteins α-SNAP_Rhg1_ and WI12_Rhg1_, which could explain the lack of enhanced resistance in *Rhg1-GmAAT* overexpression lines.

We investigated the effects of the AAT_Rhg1_ conserved site mutations Y268L and D122A on SCN resistance. We recently reported the synthetic phenotype that AAT_Rhg1_ overexpression enhances *RUBY*-mediated betalain production in soybean roots, while AAT_Rhg1_ D122A fails to do so and AAT_Rhg1_ Y268L causes even stronger pigmentation (Du *et al*., 2025). In the present work, synthesized RNAi-insensitive *SynAATRhg1WT* successfully restored SCN resistance in an iAAT soybean line. Consistent with the betalain phenotype, *SynAATRhg1D122A* failed to complement SCN resistance. The Y268 residue is predicted to reside within the internal channel of AAT_Rhg1_ near its putative substrate-binding site, and similar mutations in other amino acid transporters have been shown to alter transport activity (Errasti-Murugarren *et al*., 2019). *SynAATRhg1Y268L* did also restore SCN resistance, but unlike its stronger effect on betalain overproduction, *SynAATRhg1Y268L* did not cause greater SCN resistance than wild-type. These results suggest that although AAT_Rhg1_Y268L may enhance cytosolic tyrosine availability for betalain synthesis, this does not translate into an overt increase in SCN resistance.

AAT_Rhg1_-mediated SCN resistance may be independent of its putative amino acid transport function and instead involve other activities of the protein such as protein-protein interactions. Alternatively, Y268L might represent a more active form of AAT_Rhg1_ but substrate transport may not be limiting for SCN resistance, analogous to overexpressing wild-type AAT_Rhg1_ in an *rhg1-b* background, which also did not confer further resistance.

This study revealed that AAT_Rhg1_ protein localizes to the tonoplast in intact soybean root cells, where it likely functions as a transporter and regulates amino acid homeostasis. The addition of an N-terminal GFP tag disrupted AAT_Rhg1_ tonoplast localization and resulted in ER retention, suggesting that the N terminus may contribute to proper subcellular targeting of AAT_Rhg1_, as the first transmembrane domain in many membrane proteins often functions as a signal anchor (Wu & Hegde, 2023). In damaged cells along the SCN penetration path, AAT_Rhg1_ has also been observed in macrovesicle-like membrane structures (Han *et al*., 2023). These structures may originate from the tonoplast of cells disintegrated by SCN penetration, and although the entire cell is no longer intact, the macrovesicles could still retain biochemical or signaling functions. At the cyst nematode feeding syncytium, the large tonoplast-bounded central vacuole degenerates and the cytoplasm fills with multiple much smaller vacuole structures (Endo, 1965; Mitchum, 2016). This may partly explain the absence of pronounced AAT_Rhg1_ protein abundance elevation in the syncytium (Han *et al*., 2023), in contrast to another *Rhg1*-encoded protein, α-SNAP, which specifically accumulates in the syncytium (Bayless *et al*., 2016). The tonoplast location of AAT_Rhg1_ on a cellular structure that undergoes such dramatic remodeling during SCN parasitism may anticipate intriguing, currently unknown contributions of AAT_Rhg1_ to resistance.

Although the amino acid transporter activity of AAT_Rhg1_ has not been mechanistically demonstrated, differences in *Rhg1-GmAAT* expression influence amino acid homeostasis in soybean root cells. Yeast (*Saccharomyces cerevisiae*) *AVT6* encodes a yeast vacuolar amino acid transporter that shares structural similarities with AAT_Rhg1_ (Russnak, Konczal and McIntire, 2001; Chahomchuen *et al*., 2009) and interestingly, AVT6 was associated with a higher tyrosine production (Wu *et al*., 2020). Silencing *Rhg1-GmAAT* altered the abundance of leucine, isoleucine and tyrosine in whole-cell extracts, showing an overall trend opposite to that observed in *Rhg1-GmAAT* overexpression roots (Guo *et al*., 2019; Du *et al*., 2025). These observations suggest that precise control of *Rhg1-GmAAT* expression may be important for maintaining homeostasis of certain amino acids under both basal and stress conditions.

In *Rhg1-GmAAT*-silenced roots, genes encoding canonical biosynthetic or catabolic enzymes for leucine, isoleucine or tyrosine were not significantly differentially expressed compared with EV roots under mock-treated conditions, suggesting that 3 dpi transcript abundance changes alone do not directly explain the differences in amino acid levels. This discrepancy could reflect transient transcriptional changes not captured at our sampling time point, post-transcriptional regulation, or effects on subcellular amino acid distributions and sequestration. We quantified amino acids from whole-cell/whole-tissue extracts and it remains possible that AAT_Rhg1_ also affects subcellular or spatially restricted amino acid pools that are critical for local defense activation. The findings that AAT_Rhg1_ localizes to the tonoplast and influences amino acid homeostasis raise the possibility that altered distribution of specific amino acids between the vacuole and the cytosol, and/or vacuole-derived membrane structures at the root-nematode interface, could contribute to defense against SCN.

Plant amino acid homeostasis and amino acid transporters participate in defense signaling during biotic stress (Sonawala *et al*., 2018). A recent study has shown that microbe-associated molecular pattern (MAMP) perception induces accumulation of apoplastic amino acids, suppressing bacterial virulence in Arabidopsis (Zhang *et al*., 2023). Arabidopsis *Amino Acid Permease* (*AAP*) gene family members have been linked to susceptibility to BCN and root knot nematode (*Meloidogyne incognita*) (Elashry *et al*., 2013; Marella *et al*., 2013). Arabidopsis *Lysine Histidine Transporter 1* (*LHT1*) mediates cellular and extracellular amino acid contents and participates in regulating defense against pathogens (Liu *et al*., 2010, Rogan *et al*., 2024). Although the present study does not identify a specific mechanism by which altered amino acid levels stimulate defense, our findings with AAT_Rhg1_ are consistent with a general model that plants may sense and incorporate inappropriate amino acid levels as part of their defense signaling network.

Our transcriptomic analyses revealed that silencing *Rhg1-GmAAT* substantially alters SCN-induced gene expression profiles, impacting transcripts for multiple metabolic and defense-related pathways, any of which may be related to SCN resistance. Relative to SCN-resistant *rhg1-b* EV roots the iAAT roots, when inoculated with SCN, exhibited more extensive transcriptional responses (a greater number of DEGs). This may be related to increased nematode parasitism in plants with less resistance. Multiple MAPK signaling DEGs and especially ethylene signaling-related DEGs were upregulated more in *rhg1-b* EV than in iAAT roots, indicating that AAT_Rhg1_ influences basal defense stress signaling, possibly priming or modulating host readiness to respond to SCN infection.

The present study found down regulation of many ethylene-associated transcripts in the more SCN-susceptible iAAT line, in addition to upregulation of other ethylene-associated transcripts in the more SCN-resistant EV (*rhg1-b*) line. In Arabidopsis-BCN interactions, nematode penetration elicits an ethylene response and feeding site establishment was more successful in ethylene signaling-deficient mutants (Marhavý *et al*., 2019). Other transcriptomic studies have highlighted DEGs involved in ethylene biosynthesis and signaling at the early stages of cyst nematode infection, which may be related to SCN resistance in various soybean genetic backgrounds (Miraeiz *et al*., 2020; Chaiprom *et al*., 2024). Notably, ethylene was recently demonstrated to be a positive regulator of the expression of two *Rhg1* genes, *Rhg1-GmAAT* and *GmSNAP18*, and to contribute to SCN resistance in an *rhg1-a* background (He *et al*., 2025). Additionally, ethylene responsive transcription factors have been shown to activate the expression of downstream defense genes in response to SCN infection (Mazarei *et al*., 2002, 2007; Deng *et al*., 2024). But contrary to the above, in Arabidopsis ethylene has been implicated in beet cyst nematode attraction to roots and in nematode susceptibility (Wubben *et al*., 2001; Kammerhofer *et al*., 2015; Piya, Binder and Hewezi, 2019), and an ethylene-insensitive soybean line exhibited reduced infection by SCN (Bent *et al*., 2006). Some of the DEGs down-regulated in iAAT roots, which were more susceptible to SCN, are annotated as EIN3-binding F-box proteins whose reduced abundance could increase ethylene signaling. However, in the present study the differential impacts of *Rhg1-GmAAT* on numerous ethylene-associated transcripts overall implicate AAT_Rhg1_ as a key modulator of ethylene signaling during defense against SCN.

Overall, as is often true with RNA-seq study results, major challenges going forward are to identify and further investigate the most salient leads. The choice of leads may differ depending on if the priority is to study broader aspects of plant susceptibility to SCN rather than to identify the AAT_Rhg1_ activities that most directly modulate SCN resistance.

Metabolomics analysis showed that silencing *Rhg1-GmAAT* altered the metabolite profile in soybean roots, with a more extensive overall impact than SCN infection, suggesting this putative amino acid transporter contributes to aspects of root metabolic homeostasis beyond its role in mediating SCN resistance. Among the significantly altered mass features detected by reversed-phase negative-mode LC-MS, fatty acids, terpenoids, shikimate/phenylpropanoid compounds, and amino acid derivatives were the most represented NPC pathway categories. Within these groups, predicted flavonoids and isoflavonoids, terpenoids, and jasmonic acid related intermediates were notable due to their previously reported association with plant-nematode interactions (Dakora and Phillips, 1996; Chin, Behm and Mathesius, 2018; Guo *et al*., 2019; Meresa, Matthys and Kyndt, 2024; Jiang *et al*., 2025). These metabolomic changes offer potential biochemical context for the reduced SCN resistance observed in *Rhg1-GmAA*T-silenced plants, although the specific contributions of individual metabolites remain to be clarified. Our RNA-seq analysis did not identify consistent changes in the expression of major biosynthetic genes that would readily account for these metabolic differences, suggesting that the observed metabolite alterations may arise from mechanisms other than large transcriptional reprogramming, such as changes in substrate availability or pathway flux. Additional work will be needed to determine how AAT_Rhg1_ activity influences these metabolic outcomes.

The present study defines multiple previously unknown functional parameters of AAT_Rhg1_ including AAT_Rhg1_ subcellular location and specific impacts on pre-infection and SCN-responsive transcription and metabolic responses. Substantial work is still needed to understand the mechanisms by which AAT_Rhg1_ confers resistance. However, the present study also demonstrates that *Rhg1-GmAAT* makes significant contributions to SCN resistance against distinct SCN populations. The above insights may already assist efforts to engineer more durable SCN resistance.

## Acknowledgements

We thank Emma Nelson and Ryan Zapotocny for selection and characterization of transgenic EV and iAAT soybean lines, Ray Collier for sharing DNA resources, Senyu Chen and Andrew Scaboo for providing SCN isolates, Sarah Swanson and the Newcomb Imaging Center for confocal microscopy support and insights, and Shaojie Han for stimulating scientific discussions about AAT_Rhg1_. This work was primarily supported by Wisconsin Agricultural Experiment Station Hatch awards to A.B., with additional funding to A.B. from United Soybean Board and to H.A.M. from the USDA National Institute of Food and Agriculture, the Agriculture and Food Research Initiative competitive grant (2024-67013-42518) and the U.S. National Science Foundation grants (MCB-2404174).

## Author Contributions

Y.D. designed the research, performed research, analyzed data and wrote the paper. A.L. performed research and critically revised the paper. J.E.C. performed research, analyzed data and wrote the paper. H.M. analyzed data and critically revised the paper. A.F.B designed the research, analyzed data and wrote the paper.

## Literature cited

Acharya, K., Tande, C. and Byamukama, E. (2016) “Determination of *Heterodera glycines* Virulence Phenotypes Occurring in South Dakota,” Plant Disease, 100(11), pp. 2281–2286. Available at: 10.1094/PDIS-04-16-0572-RE.

Bandara, A.Y., Weerasooriya, D.K., Bradley, C.A., Allen, T.W. and Esker, P.D. (2020) “Dissecting the economic impact of soybean diseases in the United States over two decades,” PLOS ONE. Edited by B.B. Sahu, 15(4), p. e0231141. Available at: 10.1371/journal.pone.0231141.

Barros, F.M.D.R., Pedrinho, A., Sant’Ana, G.D.C., Freitas, C.C.G., Rosa, J.M.O., Oliveira, C.M.G.D., Rozada, C., Marto, F.N.D.S., Pascoalino, J.A.L., Silva, L.A.D. and Andreote, F.D. (2023) “Plant-parasitic nematode community and enzyme activities in soils under no-till soybean crops in Brazil,” Rhizosphere, 27, p. 100736. Available at: 10.1016/j.rhisph.2023.100736.

Bayless, A.M., Smith, J.M., Song, J., McMinn, P.H., Teillet, A., August, B.K. and Bent, A.F. (2016) “Disease resistance through impairment of α-SNAP–NSF interaction and vesicular trafficking by soybean *Rhg1*,” Proceedings of the National Academy of Sciences, 113(47). Available at: 10.1073/pnas.1610150113.

Bent, A. (2006) “*Arabidopsis thaliana* Floral Dip Transformation Method,” in W. Kan, Agrobacterium Protocols. New Jersey: Humana Press, pp. 87–104. Available at: 10.1385/1-59745-130-4:87.

Bent, A.F., Hoffman, T.K., Schmidt, J.S., Hartman, G.L., Hoffman, D.D., Ping, X., Tucker, M.L. (2006) "Disease- and Performance-Related Traits of Ethylene-Insensitive Soybean," Crop Science, 46, pp. 893–901. Available at: 10.2135/cropsci2005.08-0235.

Bent, A.F. (2022) “Exploring Soybean Resistance to Soybean Cyst Nematode,” Annual Review of Phytopathology, 60(1), pp. 379–409. Available at: 10.1146/annurev-phyto-020620-120823.

Bradley, C.A., Allen, T.W., Sisson, A.J., Bergstrom, G.C., Bissonnette, K.M., Bond, J., Byamukama, E., Chilvers, M.I., Collins, A.A., Damicone, J.P., Dorrance, A.E., Dufault, N.S., Esker, P.D., Faske, T.R., Fiorellino, N.M., Giesler, L.J., Hartman, G.L., Hollier, C.A., Isakeit, T., Jackson-Ziems, T.A., Jardine, D.J., Kelly, H.M., Kemerait, R.C., Kleczewski, N.M., Koehler, A.M., Kratochvil, R.J., Kurle, J.E., Malvick, D.K., Markell, S.G., Mathew, F.M., Mehl, H.L., Mehl, K.M., Mueller, D.S., Mueller, J.D., Nelson, B.D., Overstreet, C., Padgett, G.B., Price, P.P., Sikora, E.J., Small, I., Smith, D.L., Spurlock, T.N., Tande, C.A., Telenko, D.E.P., Tenuta, A.U., Thiessen, L.D., Warner, F., Wiebold, W.J. and Wise, K.A. (2021) “Soybean Yield Loss Estimates Due to Diseases in the United States and Ontario, Canada, from 2015 to 2019,” Plant Health Progress, 22(4), pp. 483–495. Available at: 10.1094/PHP-01-21-0013-RS.

Butler, K.J., Chen, S., Smith, J.M., Wang, X. and Bent, A.F. (2019) “Soybean Resistance Locus *Rhg1* Confers Resistance to Multiple Cyst Nematodes in Diverse Plant Species,” Phytopathology®, 109(12), pp. 2107–2115. Available at: 10.1094/PHYTO-07-19-0225-R.

Chahomchuen, T., Hondo, K., Ohsaki, M., Sekito, T. and Kakinuma, Y. (2009) “Evidence for Avt6 as a vacuolar exporter of acidic amino acids in Saccharomyces cerevisiae cells,” The Journal of General and Applied Microbiology, 55(6), pp. 409–417. Available at: 10.2323/jgam.55.409.

Chaiprom, U., Miraeiz, E., Lee, T.G., Drnevich, J. and Hudson, M. (2024) “Impact of Rhg1 copy number variation on a soybean cyst nematode resistance transcriptional network,” G3: Genes, Genomes, Genetics. Edited by A. Doust, p. jkae226. Available at: 10.1093/g3journal/jkae226.

Chen, S. (2020) “Dynamics of Population Density and Virulence Phenotype of the Soybean Cyst Nematode as Influenced by Resistance Source Sequence and Tillage,” Plant Disease, 104(8), pp. 2111–2122. Available at: 10.1094/PDIS-09-19-1916-RE.

Chin, S., Behm, C.A. and Mathesius, U. (2018) “Functions of Flavonoids in Plant– Nematode Interactions,” Plants, 7(4), p. 85. Available at: 10.3390/plants7040085.

Concibido, V.C., Diers, B.W. and Arelli, P.R. (2004) “A Decade of QTL Mapping for Cyst Nematode Resistance in Soybean,” Crop Science, 44(4), pp. 1121–1131. Available at: 10.2135/cropsci2004.1121.

Cook, D.E., Bayless, A.M., Wang, K., Guo, X., Song, Q., Jiang, J. and Bent, A.F. (2014) “Distinct Copy Number, Coding Sequence, and Locus Methylation Patterns Underlie Rhg1-Mediated Soybean Resistance to Soybean Cyst Nematode,” Plant Physiology, 165(2), pp. 630–647. Available at: 10.1104/pp.114.235952.

Cook, D.E., Lee, T.G., Guo, X., Melito, S., Wang, K., Bayless, A.M., Wang, J., Hughes, T.J., Willis, D.K., Clemente, T.E., Diers, B.W., Jiang, J., Hudson, M.E. and Bent, A.F. (2012) “Copy Number Variation of Multiple Genes at Rhg1 Mediates Nematode Resistance in Soybean,” Science, 338(6111), pp. 1206–1209. Available at: 10.1126/science.1228746.

Dakora, F.D. and Phillips, D.A. (1996) “Diverse functions of isoflavonoids in legumes transcend anti-microbial definitions of phytoalexins,” Physiological and Molecular Plant Pathology, 49(1), pp. 1–20. Available at: 10.1006/pmpp.1996.0035.

Davis, E. (2004) “Getting to the roots of parasitism by nematodes,” Trends in Parasitology, 20(3), pp. 134–141. Available at: 10.1016/j.pt.2004.01.005.

Deng, M., Zhang, L., Yang, C., Zeng, Q., Zhong, L. and Guo, X. (2024) “GmERFVII transcription factors upregulate *PATHOGENESIS-RELATED10* and contribute to soybean cyst nematode resistance,” Plant Physiology, 197(1), p. kiae548. Available at: 10.1093/plphys/kiae548.

Dong, J. and Hudson, M.E. (2022) “WI12 *Rhg1* interacts with DELLAs and mediates soybean cyst nematode resistance through hormone pathways,” Plant Biotechnology Journal, 20(2), pp. 283–296. Available at: 10.1111/pbi.13709.

Du, Y., Jung, S., Maeda, H. and Bent, A.F. (2025) “Soybean Cyst Nematode-Resistant Protein AATRhg1 Affects Amino Acid Homeostasis and Betalain Accumulation,” Plant Direct, 9(8), p. e70098. Available at: 10.1002/pld3.70098.

Dührkop, K., Fleischauer, M., Ludwig, M., Aksenov, A.A., Melnik, A.V., Meusel, M., Dorrestein, P.C., Rousu, J. and Böcker, S. (2019) “SIRIUS 4: a rapid tool for turning tandem mass spectra into metabolite structure information,” Nature Methods, 16(4), pp. 299–302. Available at: 10.1038/s41592-019-0344-8.

Elashry, A., Okumoto, S., Siddique, S., Koch, W., Kreil, D.P. and Bohlmann, H. (2013) “The AAP gene family for amino acid permeases contributes to development of the cyst nematode Heterodera schachtii in roots of Arabidopsis,” Plant Physiology and Biochemistry, 70, pp. 379–386. Available at: 10.1016/j.plaphy.2013.05.016.

El-Azaz, J. and Maeda, H.A. (2024) “A simplified liquid chromatography-mass spectrometry methodology to probe the shikimate and aromatic amino acid biosynthetic pathways in plants,” The Plant Journal, 120(5), pp. 2286–2304. Available at: 10.1111/tpj.17105.

Endo, B. Y. (1965). Histological responses of resistant and susceptible soybean varieties, and backcross progeny to entry and development of *Heterodera glycines*. Phytopathology 55:375–381

Errasti-Murugarren, E., Fort, J., Bartoccioni, P., Díaz, L., Pardon, E., Carpena, X., Espino-Guarch, M., Zorzano, A., Ziegler, C., Steyaert, J., Fernández-Recio, J., Fita, I. and Palacín, M. (2019) “L amino acid transporter structure and molecular bases for the asymmetry of substrate interaction,” Nature Communications, 10(1), p. 1807. Available at: 10.1038/s41467-019-09837-z.

Estrada-Navarrete, G. and Alvarado-Affantranger, X. (no date) “Fast, efficient and reproducible genetic transformation of Phaseolus spp. by Agrobacterium rhizogenes.”

Fan, Y. (2020) “One-step generation of composite soybean plants with transgenic roots by Agrobacterium rhizogenes-mediated transformation.”

Gardner, M., Heinz, R., Wang, J. and Mitchum, M.G. (2017) “Genetics and Adaptation of Soybean Cyst Nematode to Broad Spectrum Soybean Resistance,” G3 Genes|Genomes|Genetics, 7(3), pp. 835–841. Available at: 10.1534/g3.116.035964.

Geldner, N., Dénervaud-Tendon, V., Hyman, D.L., Mayer, U., Stierhof, Y. and Chory, J. (2009) “Rapid, combinatorial analysis of membrane compartments in intact plants with a multicolor marker set,” The Plant Journal, 59(1), pp. 169–178. Available at: 10.1111/j.1365-313X.2009.03851.x.

Guo, W., Zhang, F., Bao, A., You, Q., Li, Z., Chen, J., Cheng, Y., Zhao, W., Shen, X., Zhou, X. and Jiao, Y. (2019) “The soybean *Rhg1* amino acid transporter gene alters glutamate homeostasis and jasmonic acid-induced resistance to soybean cyst nematode,” Molecular Plant Pathology, 20(2), pp. 270–286. Available at: 10.1111/mpp.12753.

Haarith, D., Das, S., Nelson, E., Zaptocony, R. and Bent, A. (2024) “Overexpression of α- SNAPRhg1 can improve *rhg1-a* mediated soybean resistance to soybean cyst nematode.” Plant Biology. Available at: 10.1101/2024.05.18.594834.

Hallgren, J., Tsirigos, K.D., Pedersen, M.D., Almagro Armenteros, J.J., Marcatili, P., Nielsen, H., Krogh, A. and Winther, O. (2022) “DeepTMHMM predicts alpha and beta transmembrane proteins using deep neural networks.” Available at: 10.1101/2022.04.08.487609.

Han, S., Smith, J.M., Du, Y. and Bent, A.F. (2023) “Soybean transporter AAT *Rhg1* abundance increases along the nematode migration path and impacts vesiculation and ROS,” Plant Physiology, 192(1), pp. 133–153. Available at: 10.1093/plphys/kiad098.

He, L., Liu, Q., Huang, J., Cai, Y., Chen, L., Wang, Y., Wang, D., Wang, C., Bent, A. and Han, S. (2025) “The AP2/ERF transcription factor GmTINY mediates ethylene regulation of Rhg1 resistance against soybean cyst nematode,” Plant Communications, p. 101378. Available at: 10.1016/j.xplc.2025.101378.

He, Y., Zhang, T., Sun, H., Zhan, H. and Zhao, Y. (2020) “A reporter for noninvasively monitoring gene expression and plant transformation.”

Howland, A., Monnig, N., Mathesius, J., Nathan, M. and Mitchum, M.G. (2018) “Survey of *Heterodera glycines* Population Densities and Virulence Phenotypes During 2015–2016 in Missouri,” Plant Disease, 102(12), pp. 2407–2410. Available at: 10.1094/PDIS-04-18-0650-SR.

Hu, Y., You, J., Li, C., Williamson, V.M. and Wang, C. (2017) “Ethylene response pathway modulates attractiveness of plant roots to soybean cyst nematode Heterodera glycines,” Scientific Reports, 7(1), p. 41282. Available at: 10.1038/srep41282.

Hua, C., Li, C., Hu, Y., Mao, Y., You, J., Wang, M., Chen, J., Tian, Z. and Wang, C. (2018) “Identification of HG Types of Soybean Cyst Nematode *Heterodera glycines* and Resistance Screening on Soybean Genotypes in Northeast China,” Journal of Nematology, 50(1), pp. 41–50. Available at: 10.21307/jofnem-2018-007.

Ithal, N., Nettleton, D., Hearne, L., Maier, T., Baum, T.J. and Mitchum, M.G. (2007) “Parallel Genome-Wide Expression Profiling of Host and Pathogen During Soybean Cyst Nematode Infection of Soybean,” Molecular Plant-Microbe Interactions®, 20(3), pp. 293–305. Available at: 10.1094/MPMI-20-3-0293.

Ithal, N., Recknor, J., Nettleton, D., Maier, T., Baum, T.J. and Mitchum, M.G. (2007) “Developmental Transcript Profiling of Cyst Nematode Feeding Cells in Soybean Roots,” Molecular Plant-Microbe Interactions®, 20(5), pp. 510–525. Available at: 10.1094/MPMI-20-5-0510.

Jiang, H., Qu, S., Liu, F., Sun, H., Li, H., Teng, W., Zhan, Y., Li, Y., Han, Y. and Zhao, X. (2025) “Multi-omics analysis identified the *GmUGT88A1* gene, which coordinately regulates soybean resistance to cyst nematode and isoflavone content,” *Plant Biotechnology Journal*, p. pbi.14586. Available at: 10.1111/pbi.14586.

Júnior, M.T.A., Santos, A.F.D., Barros, D.I., Quirino, M.S., Nunes, H.V., Sales, P.V.G., Oliveira, L.B.D., Leite, O.D.C., Gonçalves, F.B., Reis, A.D.S., Vale, K.C.L., Pinto, J.R., Aguiar, E.C.D. and Nunes, B.H.D.N. (2022) “Nematodes, a Problem for Soybean Crops in the Brazilian Cerrado,” Journal of Scientific Research and Reports, pp. 1–6. Available at: 10.9734/jsrr/2022/v28i630522.

Juvale, P.S. and Baum, T.J. (2018) “‘Cyst-ained’ research into Heterodera parasitism,” PLOS Pathogens. Edited by C. Zipfel, 14(2), p. e1006791. Available at: 10.1371/journal.ppat.1006791.

Kammerhofer, N., Radakovic, Z., Regis, J.M.A., Dobrev, P., Vankova, R., Grundler, F.M.W., Siddique, S., Hofmann, J. and Wieczorek, K. (2015) “Role of stress-related hormones in plant defence during early infection of the cyst nematode *Heterodera schachtii* in Arabidopsis,” New Phytologist, 207(3), pp. 778–789. Available at: 10.1111/nph.13395.

Kandoth, P.K., Ithal, N., Recknor, J., Maier, T., Nettleton, D., Baum, T.J. and Mitchum, M.G. (2011) “The Soybean *Rhg1* Locus for Resistance to the Soybean Cyst Nematode *Heterodera glycines* Regulates the Expression of a Large Number of Stress- and Defense- Related Genes in Degenerating Feeding Cells,” Plant Physiology, 155(4), pp. 1960–1975. Available at: 10.1104/pp.110.167536.

Kim, H.W., Wang, M., Leber, C.A., Nothias, L.-F., Reher, R., Kang, K.B., van der Hooft, J.J.J., Dorrestein, P.C., Gerwick, W.H. and Cottrell, G.W. (2021) “NPClassifier: A Deep Neural Network-Based Structural Classification Tool for Natural Products,” Journal of Natural Products, 84(11), pp. 2795–2807. Available at: 10.1021/acs.jnatprod.1c00399.

Kim, Y.-H., Kim, K.-S. and Riggs, R.D. (2010) “Differential Subcellular Responses in Resistance Soybeans Infected with Soybean Cyst Nematode Races,” The Plant Pathology Journal, 26(2), pp. 154–158. Available at: 10.5423/PPJ.2010.26.2.154.

Kim, Y.H., Kim, K.S. and Riggs, R.D. (2012) “Initial Subcellular Responses of Susceptible and Resistant Soybeans Infected with the Soybean Cyst Nematode,” The Plant Pathology Journal, 28(4), pp. 401–408. Available at: 10.5423/PPJ.OA.04.2012.0054.

Klink, V.P., Hosseini, P., Matsye, P.D., Alkharouf, N.W. and Matthews, B.F. (2010) “Syncytium gene expression in Glycine max[PI 88788] roots undergoing a resistant reaction to the parasitic nematode Heterodera glycines,” Plant Physiology and Biochemistry, 48(2– 3), pp. 176–193. Available at: 10.1016/j.plaphy.2009.12.003.

Lee, T.G., Kumar, I., Diers, B.W. and Hudson, M.E. (2015) “Evolution and selection of *R hg1*, a copy-number variant nematode-resistance locus,” Molecular Ecology, 24(8), pp. 1774–1791. Available at: 10.1111/mec.13138.

Liu, G., Ji, Y., Bhuiyan, N.H., Pilot, G., Selvaraj, G., Zou, J. and Wei, Y. (2010) “Amino Acid Homeostasis Modulates Salicylic Acid–Associated Redox Status and Defense Responses in *Arabidopsis*,” The Plant Cell, 22(11), pp. 3845–3863. Available at: 10.1105/tpc.110.079392.

Liu, S., Kandoth, P.K., Lakhssassi, N., Kang, J., Colantonio, V., Heinz, R., Yeckel, G., Zhou, Z., Bekal, S., Dapprich, J., Rotter, B., Cianzio, S., Mitchum, M.G. and Meksem, K. (2017) “The soybean GmSNAP18 gene underlies two types of resistance to soybean cyst nematode,” Nature Communications, 8(1), p. 14822. Available at: 10.1038/ncomms14822.

Marella, H.H., Nielsen, E., Schachtman, D.P. and Taylor, C.G. (2013) “The Amino Acid Permeases AAP3 and AAP6 Are Involved in Root-Knot Nematode Parasitism of *Arabidopsis*,” Molecular Plant-Microbe Interactions®, 26(1), pp. 44–54. Available at: 10.1094/MPMI-05-12-0123-FI.

Marhavý, P., Kurenda, A., Siddique, S., Dénervaud Tendon, V., Zhou, F., Holbein, J., Hasan, M.S., Grundler, F.M., Farmer, E.E. and Geldner, N. (2019) “Single-cell damage elicits regional, nematode-restricting ethylene responses in roots,” The EMBO Journal, 38(10), p. e100972. Available at: 10.15252/embj.2018100972.

Mazarei, M., Elling, A.A., Maier, T.R., Puthoff, D.P. and Baum, T.J. (2007) “GmEREBP1 Is a Transcription Factor Activating Defense Genes in Soybean and *Arabidopsis*,” Molecular Plant-Microbe Interactions®, 20(2), pp. 107–119. Available at: 10.1094/MPMI-20-2-0107.

Mazarei, M., Puthoff, D.P., Hart, J.K., Rodermel, S.R. and Baum, T.J. (2002) “Identification and Characterization of a Soybean Ethylene-Responsive Element-Binding Protein Gene Whose mRNA Expression Changes During Soybean Cyst Nematode Infection,” Molecular Plant-Microbe Interactions®, 15(6), pp. 577–586. Available at: 10.1094/MPMI.2002.15.6.577.

McCarville, M.T., Marett, C.C., Mullaney, M.P., Gebhart, G.D. and Tylka, G.L. (2017) “Increase in Soybean Cyst Nematode Virulence and Reproduction on Resistant Soybean Varieties in Iowa From 2001 to 2015 and the Effects on Soybean Yields,” Plant Health Progress, 18(3), pp. 146–155. Available at: 10.1094/PHP-RS-16-0062.

Meresa, B.K., Matthys, J. and Kyndt, T. (2024) “Biochemical Defence of Plants against Parasitic Nematodes,” Plants, 13(19), p. 2813. Available at: 10.3390/plants13192813.

Miraeiz, E., Chaiprom, U., Afsharifar, A., Karegar, A., M. Drnevich, J. and E. Hudson, M. (2020) “Early transcriptional responses to soybean cyst nematode HG Type 0 show genetic differences among resistant and susceptible soybeans,” Theoretical and Applied Genetics, 133(1), pp. 87–102. Available at: 10.1007/s00122-019-03442-w.

Mitchum, M.G. (2016) “Soybean Resistance to the Soybean Cyst Nematode *Heterodera glycines* : An Update,” Phytopathology®, 106(12), pp. 1444–1450. Available at: 10.1094/PHYTO-06-16-0227-RVW.

Niblack, T. L., Arelli, P. R., Noel, G. R., Opperman, C. H., Orf, J. H., Schmitt, D. P., Shannon, J. G., & Tylka, G. L. (2002). A Revised Classification Scheme for Genetically Diverse Populations of *Heterodera glycines*. Journal of nematology, 34(4), 279–288.

Ødum, M.T., Teufel, F., Thumuluri, V., Almagro Armenteros, J.J., Johansen, A.R., Winther, O. and Nielsen, H. (2024) “DeepLoc 2.1: multi-label membrane protein type prediction using protein language models,” Nucleic Acids Research, 52(W1), pp. W215–W220. Available at: 10.1093/nar/gkae237.

Patil, G.B., Lakhssassi, N., Wan, J., Song, L., Zhou, Z., Klepadlo, M., Vuong, T.D., Stec, A.O., Kahil, S.S., Colantonio, V., Valliyodan, B., Rice, J.H., Piya, S., Hewezi, T., Stupar, R.M., Meksem, K. and Nguyen, H.T. (2019) “Whole-genome re-sequencing reveals the impact of the interaction of copy number variants of the *rhg1* and *Rhg4* genes on broad-based resistance to soybean cyst nematode,” Plant Biotechnology Journal, 17(8), pp. 1595–1611. Available at: 10.1111/pbi.13086.

Paysan-Lafosse, T., Blum, M., Chuguransky, S., Grego, T., Pinto, B.L., Salazar, G.A., Bileschi, M.L., Bork, P., Bridge, A., Colwell, L., Gough, J., Haft, D.H., Letunić, I., Marchler- Bauer, A., Mi, H., Natale, D.A., Orengo, C.A., Pandurangan, A.P., Rivoire, C., Sigrist, C.J.A., Sillitoe, I., Thanki, N., Thomas, P.D., Tosatto, S.C.E., Wu, C.H. and Bateman, A. (2023) “InterPro in 2022,” Nucleic Acids Research, 51(D1), pp. D418–D427. Available at: 10.1093/nar/gkac993.

Peng, D., Jiang, R., Peng, H. and Liu, S. (2021) “Soybean cyst nematodes: a destructive threat to soybean production in China,” Phytopathology Research, 3(1), p. 19. Available at: 10.1186/s42483-021-00095-w.

Piya, S., Binder, B.M. and Hewezi, T. (2019) “Canonical and noncanonical ethylene signaling pathways that regulate Arabidopsis susceptibility to the cyst nematode *Heterodera schachtii*,” New Phytologist, 221(2), pp. 946–959. Available at: 10.1111/nph.15400.

Rogan, C.J., Pang, Y.-Y., Mathews, S.D., Turner, S.E., Weisberg, A.J., Lehmann, S., Rentsch, D. and Anderson, J.C. (2024) “Transporter-mediated depletion of extracellular proline directly contributes to plant pattern-triggered immunity against a bacterial pathogen,” Nature Communications, 15(1), p. 7048. Available at: 10.1038/s41467-024-51244-6.

Russnak, R., Konczal, D. and McIntire, S.L. (2001) “A Family of Yeast Proteins Mediating Bidirectional Vacuolar Amino Acid Transport,” Journal of Biological Chemistry, 276(26), pp. 23849–23857. Available at: 10.1074/jbc.M008028200.

Schmid, R., Heuckeroth, S., Korf, A., Smirnov, A., Myers, O., Dyrlund, T.S., Bushuiev, R., Murray, K.J., Hoffmann, N., Lu, M., Sarvepalli, A., Zhang, Z., Fleischauer, M., Dührkop, K., Wesner, M., Hoogstra, S.J., Rudt, E., Mokshyna, O., Brungs, C., Ponomarov, K., Mutabdžija, L., Damiani, T., Pudney, C.J., Earll, M., Helmer, P.O., Fallon, T.R., Schulze, T., Rivas-Ubach, A., Bilbao, A., Richter, H., Nothias, L.-F., Wang, M., Orešič, M., Weng, J.-K., Böcker, S., Jeibmann, A., Hayen, H., Karst, U., Dorrestein, P.C., Petras, D., Du, X. and Pluskal, T. (2023) “Integrative analysis of multimodal mass spectrometry data in MZmine 3,” Nature Biotechnology, 41(4), pp. 447–449. Available at: 10.1038/s41587-023-01690-2.

Shi, X., Chen, Q., Liu, S., Wang, J., Peng, D. and Kong, L. (2021) “Combining targeted metabolite analyses and transcriptomics to reveal the specific chemical composition and associated genes in the incompatible soybean variety PI437654 infected with soybean cyst nematode HG1.2.3.5.7,” BMC Plant Biology, 21(1), p. 217. Available at: 10.1186/s12870-021-02998-4.

Sonawala, U., Dinkeloo, K., Danna, C.H., McDowell, J.M. and Pilot, G. (2018) “Review: Functional linkages between amino acid transporters and plant responses to pathogens,” Plant Science, 277, pp. 79–88. Available at: 10.1016/j.plantsci.2018.09.009.

Song, W., Qi, N., Liang, C., Duan, F. and Zhao, H. (2019) “Soybean root transcriptome profiling reveals a nonhost resistant response during Heterodera glycines infection,” PLOS ONE. Edited by K. Ulaganathan, 14(5), p. e0217130. Available at: 10.1371/journal.pone.0217130.

Sultana, M.S., Niyikiza, D., Hawk, T.E., Coffey, N., Lopes-Caitar, V., Pfotenhauer, A.C., El- Messidi, H., Wyman, C., Pantalone, V. and Hewezi, T. (2024) “Differential transcriptome reprogramming induced by the soybean cyst nematode Type 0 and Type 1.2.5.7 during resistant and susceptible interactions,” *Molecular Plant-Microbe Interactions®*, p. MPMI- 08–24-0092-R. Available at: 10.1094/MPMI-08-24-0092-R.

Torabi, S., Seifi, S., Geddes-McAlister, J., Tenuta, A., Wally, O., Torkamaneh, D. and Eskandari, M. (2023) “Soybean–SCN Battle: Novel Insight into Soybean’s Defense Strategies against Heterodera glycines,” International Journal of Molecular Sciences, 24(22), p. 16232. Available at: 10.3390/ijms242216232.

Tylka, G.L. and Marett, C.C. (2021) “Known Distribution of the Soybean Cyst Nematode, *Heterodera glycines* , in the United States and Canada in 2020,” Plant Health Progress, 22(1), pp. 72–74. Available at: 10.1094/PHP-10-20-0094-BR.

Weber, E., Engler, C., Gruetzner, R., Werner, S. and Marillonnet, S. (2011) “A Modular Cloning System for Standardized Assembly of Multigene Constructs,” PLoS ONE. Edited by J. Peccoud, 6(2), p. e16765. Available at: 10.1371/journal.pone.0016765.

Wei, H., Lian, Y., Li, J., Li, H., Song, Q., Wu, Y., Lei, C., Wang, S., Zhang, H., Wang, J. and Lu, W. (2022) “Identification of Candidate Genes Controlling Soybean Cyst Nematode Resistance in ‘Handou 10’ Based on Genome and Transcriptome Analyzes,” Frontiers in Plant Science, 13, p. 860034. Available at: 10.3389/fpls.2022.860034.

Wu, H. and Hegde, R.S. (2023) “Mechanism of signal-anchor triage during early steps of membrane protein insertion,” Molecular Cell, 83(6), pp. 961–973.e7. Available at: 10.1016/j.molcel.2023.01.018.

Wu, Y., Cai, M., Song, X., Li, Y., Wang, H., Mao, J., Liu, Q., Xu, H. and Qiao, M. (2020) “Comparative transcriptome analysis of genomic region deletion strain with enhanced l- tyrosine production in Saccharomyces cerevisiae,” Biotechnology Letters, 42(3), pp. 453–460. Available at: 10.1007/s10529-019-02784-1.

Wubben, M.J.E., Su, H., Rodermel, S.R. and Baum, T.J. (2001) “Susceptibility to the Sugar Beet Cyst Nematode Is Modulated by Ethylene Signal Transduction in *Arabidopsis thaliana*,” Molecular Plant-Microbe Interactions®, 14(10), pp. 1206–1212. Available at: 10.1094/MPMI.2001.14.10.1206.

Zhang, L., Lilley, C.J., Imren, M., Knox, J.P. and Urwin, P.E. (2017) “The Complex Cell Wall Composition of Syncytia Induced by Plant Parasitic Cyst Nematodes Reflects Both Function and Host Plant,” Frontiers in Plant Science, 8, p. 1087. Available at: 10.3389/fpls.2017.01087.

Zhang, L., Zeng, Q., Zhu, Q., Tan, Y. and Guo, X. (2022) “Essential Roles of Cupredoxin Family Proteins in Soybean Cyst Nematode Resistance,” Phytopathology®, 112(7), pp. 1545–1558. Available at: 10.1094/PHYTO-09-21-0391-R.

Zhang, X., Tubergen, P.J., Agorsor, I.D.K., Khadka, P., Tembe, C., Denbow, C., Collakova, E., Pilot, G. and Danna, C.H. (2023) “Elicitor-induced plant immunity relies on amino acids accumulation to delay the onset of bacterial virulence,” Plant Physiology, 192(1), pp. 601– 615. Available at: 10.1093/plphys/kiad048.

