## Supplemental Materials and Methods for "AAT_Rhg1_ is a tonoplast protein that alters amino acid, metabolic and defense responses and nematode resistance"

### Extended Materials and Methods Descriptions

### Plasmid constructs

The *Rhg1-GmAAT* silencing construct was described in (Du *et al*., 2025). In brief, the hairpin RNA construct was generated using a 292 bp DNA fragment spanning the junction between *Rhg1-GmAAT* first and second exons, followed by a 523bp GUS fragment as a spacer and then the same 292bp fragment in reverse complement orientation, cloned into pCambia2300 plasmid under the control of a *GmUbi* promoter (Hernandez-Garcia *et al*., 2010) and an *Rbcs-E9* terminator. A spectinomycin resistance marker with a chloroplast targeting signal driven by a double Cauliflower mosaic virus (CaMV) *35S* promoter and nopaline synthase (*nos*) terminator was used for selecting whole transgenic soybeans (Martinell *et al*., 2013).

An artificial RNAi-escaping *Rhg1-GmAAT* coding sequence (CDS) was created by designing synonymous codon substitutions based on soybean codon usage table. A DNA fragment containing the designed RNAi-escaping *Rhg1-GmAAT* CDS with BbsI cutting sites flanking the CDS was synthesized at Twist Bioscience (South San Francisco, CA, USA) and cloned into a Golden Gate level 0 and then level 1 vector under control of a *GmUbi* promoter and *nos* terminator. The *Rhg1-GmAAT* expression module together with *2xCaMV* $35S_{pro}$*:TMV omega enhancer:ZsGreen-KDEL:*$NOS_{ter}$ were cloned into binary vector pAGM4673, with the *ZsGreen* module next to the left border of the T-DNA as the screenable marker for transgenic soybean roots. Single amino acid mutation *Rhg1-GmAAT* alleles were generated by site-directed mutagenesis by PCR with overlapping oligonucleotide primers, followed by *DpnI* digestion of template DNA and transformation into *E. coli*.

To investigate the effect of elevated *Rhg1-GmAAT* transcript levels on SCN resistance, two T-DNA cassettes were constructed and inserted separately into binary vector pAGM4673, each along with a spectinomycin resistance marker gene expression cassette for selection of transgenic plants. The native *Rhg1-GmAAT* construct was built from the Williams82 *Glyma.18G022400* genomic sequence, including the full 1,546 bp native promoter spanning the region upstream of the 5' untranslated region (UTR) to the transcription start of divergently transcribed *Glyma.18G022500*, and also the complete coding sequence including introns and UTRs followed by the 440 bp downstream region serving as the native terminator. The overexpression construct consisted of the *Rhg1-GmAAT* coding sequence (exons only) controlled by a double CaMV *35S* promoter with a TMV *omega* enhancer, followed by the *nos* terminator.

For microscopy, genes encoding the fluorescent proteins mCherry, TdTomato with C-terminal KDEL endoplasmic reticulum (ER) retention signal, and mWasabi were obtained as Golden Gate level 0 plasmids. Subsequent cloning was done using the Golden Gate Assembly MoClo Tool Kit (Weber *et al.*, 2011). Green fluorescent protein mWasabi was tagged to the N or C terminus of $AAT_{Rhg1}$ in Golden Gate level 0 to level 1 reactions. The loop-tagged $AAT_{Rhg1}$ was created through Gibson Assembly. PCR reactions were performed to amplify *Rhg1-GmAAT* and *mWasabi* CDS with compatible overhangs added for Gibson Assembly. *VAMP711* was cloned from Arabidopsis Biological Resource Center (ABRC) CD3-781601, and inserted into Golden Gate level 0 vector through BbsI overhangs added by PCR. The above genes were cloned into Golden Gate level 1 vectors for expression under the control of double CaMV *35S* promoter with TMV *omega* translational enhancer and *nos* terminator. The level 1 modules were assembled into the Golden Gate level 2 binary vector to generate constructs for co-expression in soybean roots.

A list of the oligonucleotide primers used is provided as Supplemental Table S1.

**Transgenic plant materials**

IL3025N (synonyms: Illini 3025N, LD11-2170; *rhg1-b* SCN resistance haplotype) and IL3849N (synonyms: Illini 3849N, LD07-3395bf; *rhg1-a* + *Rhg4* SCN resistance haplotypes) were developed by Brian Diers, University of Illinois at Urbana-Champaign, and are available through Baird Seed Co. (Williamsfield, Illinois). The pedigree for IL3025N is Syngenta_03JR313108 x LD05-3171. IL3849N is an SCN-resistant reselection from LD07-3395 whose pedigree is Syngenta_WW115926 x LD00-2817. Whole transgenic *Rhg1-GmAAT*-silenced soybean plants were generated from IL3025N at Wisconsin Crop Innovation Center (WCIC) as described in Du *et al.*, 2025. Transgenic lines were selected using the spectinomycin resistance marker (Martinell *et al.*, 2013) and T-DNA copy numbers were determined by qPCR using genomic DNA (more details in “DNA extraction and copy number test” section). Transgene expression or target gene silencing were verified by RT-qPCR. For the *Rhg1-GmAAT*-silenced RNAi and corresponding EV lines, stable homozygous transgenic lines were selected, and 4th generation (T4) transgenic plants were used in SCN assays. For *Rhg1-GmAAT* overexpression lines generated in Williams 82 (Wm82) background, homozygous T2 plants were used in SCN assays. For *Rhg1-GmAAT* overexpression lines generated in IL3025N and IL3849N, T1 plants were used in SCN assays. Because the T1 siblings from the same event segregated and are a mixture of homozygotes, hemizygotes and non-transgenic, T-DNA copy number was tested for each plant used in SCN assays.

Composite soybean plants with transgenic roots were generated from soybean seedling hypocotyls using *Agrobacterium rhizogenes* strain K599 following the methods of Estrada-Navarrete & Alvarado-Affantranger (2007) and Fan *et al*. (2020). Detached transgenic soybean roots were produced from soybean cotyledons using the method described in Cook *et al*. (2012).

Transgenic Arabidopsis lines expressing *Rhg1-GmAAT* were generated using the floral dip method (Bent, 2006) and homozygous transgenic lines were identified using the spectinomycin resistance gene described above (Martinell *et al*., 2013). *AtAVT6C* knockout line CS864997 was ordered from ABRC (Arabidopsis Biological Resource Center, Ohio State University, Columbus OH).

**DNA extraction and copy number test**

DNA was extracted from young soybean leaf tissue after flash-freezing in liquid nitrogen, using the CTAB/chloroform method. Transgene copy number was tested by qPCR, measuring the relative level of the spectinomycin resistance gene on T-DNA relative to *Glyma.18G022800*, which is known to only have one copy per haploid genome.

**SCN propagation and resistance assay**

SCN populations were initially collected in the field and propagated in the laboratory on soybean plants grown in a 1:1 soil sand mixture. The HG 0 population used in experiments was originally collected in Illinois. Two HG 2.5.7 populations were sourced from Minnesota (HG 2.5.7 MN) or Missouri (HG 2.5.7 MO). To maintain or enhance virulence, these SCN populations were propagated on different soybean varieties: HG 0 on the susceptible variety Wm82 and HG 2.5.7 on *rhg1-b* variety IL3025N. Propagation pots were grown for approximately two months before harvesting. SCN cysts and eggs were extracted using the method described in Cool *et al*., 2012. Eggs were hatched in 4 mM pH 5.6 $ZnCl_2$ solution.

SCN resistance assays were conducted in a controlled temperature growth room set for 16-hour light (~800 μmol/m$^2$/sec) at 28$^o$C. Each transgenic plant was planted individually in 6cm pots containing a 1:1 soil-sand mixture and inoculated with nematodes. Whole transgenic plants were each inoculated with ~1000 SCN eggs and incubated for 35 days, while composite plants were each inoculated with ~750 freshly hatched SCN J2s and incubated for 30 days. Felt strips were placed at the bottom of each pot to wick water. Soybean plants with different genotypes within a single experiment were anonymized with random numbering and arranged in a completely randomized design. After the incubation period, cysts were extracted from soil by water flotation, then collected on nested U.S. Standard No. 25 (710 μm opening) and No. 80 (180 μm opening) sieves, followed by separation using 68.1% (w/v) sucrose solution. Cysts harvested from each plant were counted using a stereo dissecting microscope.

A sugar beet cyst nematode (*Heterodera schachtii*, BCN) population was obtained from Dr. Thomas J Baum and maintained on collard. BCN assays on Arabidopsis were conducted as described by Butler *et al*. (2019).

**RNA extraction, RT-qPCR, and RNA-seq**

Sampled plant tissues were flash-frozen in liquid nitrogen and stored at -80°C until RNA extraction. Frozen samples were mechanically ground with 2mm zirconium silicate bead and PowerLyzer™ 24 (MO BIO Laboratories, Inc., Carlsbad, CA, USA). RNA was extracted using the Direct-zol™ RNA MiniPrep Plus Kit (Zymo Research, Irvine, CA, USA) according to the manufacturer's protocol. To obtain high-quality RNA for RNA-seq, an additional chloroform phase separation step was included, adapted from the TRIzol™ Reagent user guide (Thermo Fisher Scientific, Waltham, MA, USA). Specifically, 1 mL of TRIzol™ was added to each sample, followed by the addition of 200 µL of chloroform. The tubes were then centrifuged at 12,000 x g for 15 minutes at 4°C. The aqueous phase was transferred to a new tube, mixed with an equal volume of 100% ethanol, and processed through the Direct-zol™ column. The extraction was then completed following Direct-zol™ manufacturer's instructions. Complementary DNA (cDNA) was synthesized using FIREScript® RT cDNA synthesis MIX from Solis BioDyne (Tartu, Estonia). Quantitative reverse transcription polymerase chain reaction (RT-qPCR) was performed using HOT FIREPol® EvaGreen® qPCR Supermix from Solis BioDyne and CFX96 real-time PCR detection system (BioRad, Hercules, CA, USA).

For RNA-seq studies, transgenic soybean plants were germinated in vermiculite for 4 days before being transplanted to sand individually. Ten days after germination, each plant was inoculated with 1000 freshly hatched SCN (HG 0) J2s or the same volume of SCN-free hatching solution for the mock treatment. Three days after inoculation, root segments with visible lesions caused by nematode penetration were collected from SCN-inoculated plants by freezing in liquid nitrogen within one minute of excision, and root segments from corresponding regions on mock-inoculated plants were sampled. Four replicate samples were analyzed for each genotype-treatment. Each replicate was comprised of multiple root sections from one individual plant. RNA-seq and data analysis were performed by Novogene (Sacramento, CA, USA) using their Eukaryotic mRNA-seq service (Illumina NovaSeq platform, 150bp paired-end reads, ≥ 20 million read pairs per sample, read quality control, and gene expression quantification). The iAAT_Mock group had three replicates after one outlier was removed. Differential expression profiling primarily utilized Novogene platform for fpkm, log2 fold change and adjusted p value ($p_{adj}$) data, as well as KEGG and GO term functional group enrichment analysis.

**Metabolite extraction and quantification**

For metabolite extraction, seeds were initially germinated in vermiculite and transferred to sand four days post-germination. At 12 days post-germination, each plant was either inoculated with 1,000 SCN J2 juveniles or mock-treated. Three days post-inoculation (dpi), root infection zones under SCN (HG 0) infection or corresponding regions from mock-treated plants were collected. For each sample, root sections from three individual plants were pooled, flash-frozen in liquid nitrogen, and freeze-dried for

48 hours using the FreeZone® 4.5L Benchtop Freeze Dryer (Labconco, Kansas City, MO, USA).

Metabolites were extracted using a 2:1 mix of methanol and chloroform, with 50 μM of $^{13}C_6$-ring-labeled-L-phenylalanine as internal standard. Approximately 50 mg of freeze-dried root tissue were added to 800 μL of extraction solvent, then vortexed continuously at ambient temperature for 45 minutes. After spinning at 20,000 *g* for five min, the supernatant (~800 μL) was transferred to a clean tube and mixed with 600 μL of LC-MS grade water and 300 μL of chloroform by vortexing for ~one minute. Following a second centrifugation step at 20,000 *g* for five min, the top aqueous layer (~1 mL) was recovered and evaporated overnight in a SpeedVac at room temperature. Dried extracts were resuspended into 50 μL of 80% methanol by vortexing and sonication in a water bath for five min, followed by centrifugation at at 20,000 *g* for five min. The final supernatant was collected and transferred to vials for LC-MS analysis.

For LC-MS analysis, a Vanquish Horizon Binary UHPLC (Thermo Scientific, Waltham, MA, USA) coupled to a Q Exactive Orbitrap mass spectrometer (Thermo Scientific, Waltham, MA, USA), was used. Amino acids were analyzed by zwitterionic hydrophilic interaction liquid chromatography (HILIC-Z) using an InfinityLab Poroshell 120 HILIC-Z column (2.7 μm particle size, 150 × 2.1 mm) (Agilent, Santa Clara, CA, USA) in a 22 min gradient of A (5 mM ammonium acetate, 0.2% acetic acid in water) in B (5 mM ammonium acetate, 0.2% acetic acid in 95% acetonitrile) at a flow rate of 0.45 mL/min: 0 to 1 min, isocratic 100% B; 1 to 7.5 min, 100% to 82% B; 7.5 to 11 min, 82 to 63% B; 11 to 14 min, 63% to 15% B; 14 to 16.5 min, isocratic 15% B; 16.5 to 17.5 min, 15% to 100% B; 17.5 to 22 min, isocratic 100% B. Total ion current (TIC). MS1 and MS2 data were collected between 1 and 16.5 minutes in full scan mode between *m/z* 70 to 1050 in positive ionization. MS/MS settings for HILIC-Z chromatography were described in El-Azaz & Maeda, 2024. Phenylpropanoids and other semi-polar metabolites were determined by reversed-phase (RP) chromatography using an Acquity UPLC HSS T3 column (1.8 µm particle size, 2.1 mm X 100 mm) (Waters, Milford, MA, USA) using two alternative gradients of A (0.1% formic acid in water) in B (0.1% formic acid in 90% acetonitrile) depending of the ionization mode (negative or positive, see below). The following 30 min gradient at 0.4 mL/min was used in negative ionization mode: 0 to 1 min, isocratic 1% B; 1 to 5 min, 1% to 10% B; 5 to 20 min, 10% to 25% B; 20 to 22 min, 25% to 99% B; 22 to 25 min, isocratic 99% B; 25 to 26 min, 100% to 1% B; 26 to 30 min, isocratic 1% B. For detection of phenylpropanoids and other semi-polar metabolites in positive ionization mode, the following 27 min gradient at 0.4 mL/min was used: 0 to 1 min, isocratic 1% B; 1 to 10 min, 1% to 10% B; 10 to 13 min, 10% to 25% B; 13 to 18 min, 25% to 99% B; 18 to 22 min, isocratic 99% B; 22 to 23.5 min, 99% to 1% B; 23.5 to 27 min, isocratic 1% B. MS1 and MS2 data were collected between 0.5 and 25 min (for the negative ionization method) and between 0.5 to 22 min (for the positive ionization method) using full scan mode between m/z 60 to 900. All other MS/MS settings for RP chromatography were described in Du *et al.*, 2025. Untargeted metabolite data processing and metabolite annotation were performed using MZmine v4.0 (Schmid *et al.*, 2023) and SIRIUS v5 (Dührkop *et al.*, 2019) as described in El-Azaz and Maeda, 2024. For targeted metabolite analysis, compound abundance was calculated based on the peak area of authentic standards and normalized to sample mass and internal standard recovery for each sample.

### Confocal microscopy

Confocal images were acquired using a Zeiss LSM 980 with 40x water immersion objective lens. TdTomato or mCherry signal was acquired with 561 nm laser excitation and 573 to 628 nm emission, while mWasabi signal was acquired with 488 nm laser excitation and 500 to 553 nm emission with beam splitter MBS 488/561 and laser intensity 2.0%.

### Statistical analyses

RNA-seq analyses are described separately. All other statistical analyses, including principal component analysis, were performed in R (version 4.4.2). Differences between two groups were evaluated using Student's *t*-test when data met assumptions of normality, or the Wilcoxon rank-sum test otherwise. For comparisons among more than two groups, one-way or two-way analysis of variance (ANOVA) followed by Tukey's HSD post hoc test was used. A $p$-value $< 0.05$ was considered statistically significant.
