## Supplemental Figures for "AAT_Rhg1_ is a tonoplast protein that alters amino acid, metabolic and defense responses and nematode resistance"

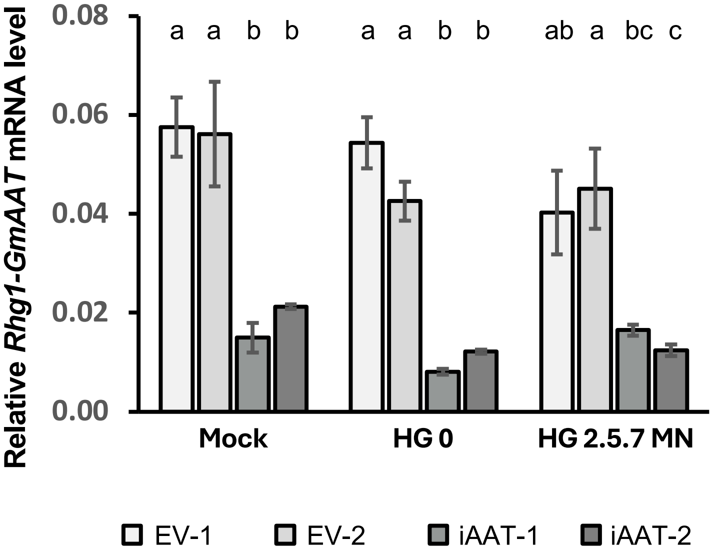


**Supplemental Figure S1. *Rhg1-GmAAT* is effectively silenced in RNAi transgenic soybean lines.**
Transcript levels of *Rhg1-GmAAT* relative to *GmNREG* in IL3025N (*rhg1-b*) transgenic lines transformed with either an empty vector (EV) or an *Rhg1-GmAAT* RNAi-silenced (iAAT) construct, following mock treatment or inoculation with SCN HG 0 or HG 2.5.7 from Minnesota (HG 2.5.7 MN) at 3 days post-inoculation (dpi). Data were analyzed using ANOVA with post-hoc Tukey HSD (n = 3). Bars sharing the same letter within each SCN treatment are not significantly different. Data are presented as mean ± SE.

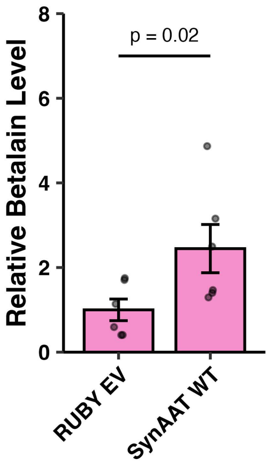


**Supplemental Figure S2. Synthesized codon-substituted RNAi-escaping *Rhg1-GmAAT* functions in *RUBY*-mediated betalain synthesis.**
Synthesized codon-substituted RNAi-escaping *Rhg1-GmAAT* wild-type (*SynAAT WT*) was co-expressed with *RUBY* in Williams 82 roots. Betalains were quantified as ($A_{538}$ + $A_{476}$) per gram of fresh weight, and normalized to roots expressing *RUBY*-alone empty vector (*RUBY* EV). Data were analyzed using one-tailed *t*-test and presented as mean ± SE. Each dot represents one sample.

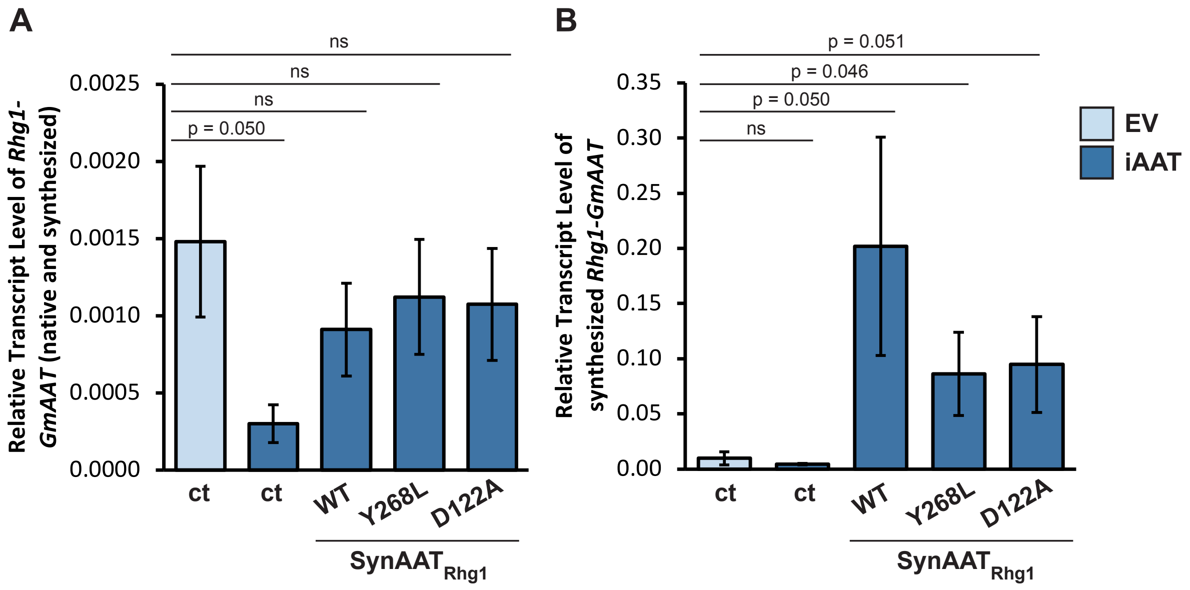


**Supplemental Figure S3. Transcript levels of *SynAAT$_{Rhg1}$* in soybean roots.** Transgenic EV plants further transformed with a marker-only construct (ct) were used as a high-expression control, and transgenic iAAT plants further transformed with a marker-only construct (ct) were used as a low-expression control. Transcript levels of *SynAAT$_{Rhg1}$* wild type (WT), *SynAAT$_{Rhg1}$ Y268L*, or *SynAAT$_{Rhg1}$ D122A* were measured using primers amplifying both native and synthetic *Rhg1-GmAAT* **(A)** or only synthetic *SynAAT$_{Rhg1}$* **(B)**. Data were analyzed using one-tailed *t*-test and are presented as mean ± SE. n = 3 or 4.

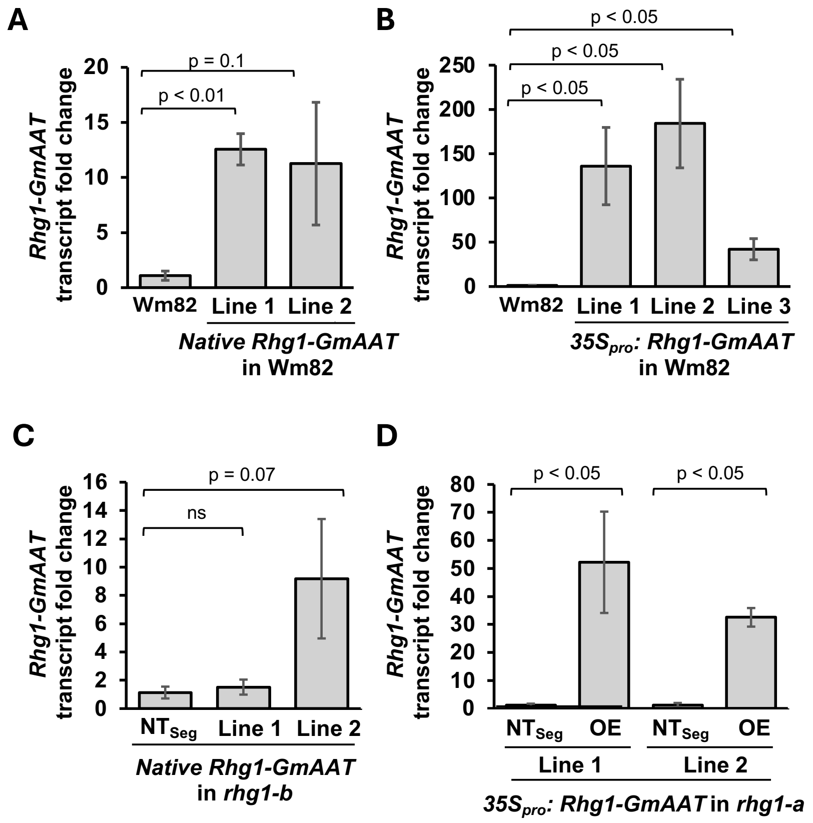


**Supplemental Figure S4. Overexpression of *Rhg1-GmAAT* in transgenic soybean lines.**

Transcript levels of *Rhg1-GmAAT* relative to *GmNREG* in the following transgenic soybean lines:

**(A)** Williams 82 transformed with a native *Rhg1-GmAAT* construct (native promoter, UTRs, exons, introns and terminator)

**(B)** Williams 82 transformed with a *35S promoter:Rhg1-GmAAT CDS:Nos terminator* construct.

**(C)** IL3025N (*rhg1-b*) transformed with the native *Rhg1-GmAAT* construct. Non-transgenic progeny from transformed plants ($NT_{seg}$) were used as controls.

**(D)** IL3849N (*rhg1-a*) transformed with the *35S* overexpression construct. $NT_{seg}$ plants were used as controls.

Data were analyzed using one-tailed *t*-test (n = 3) comparing transgenic lines to non-transgenic controls, and are presented as mean ± SE.

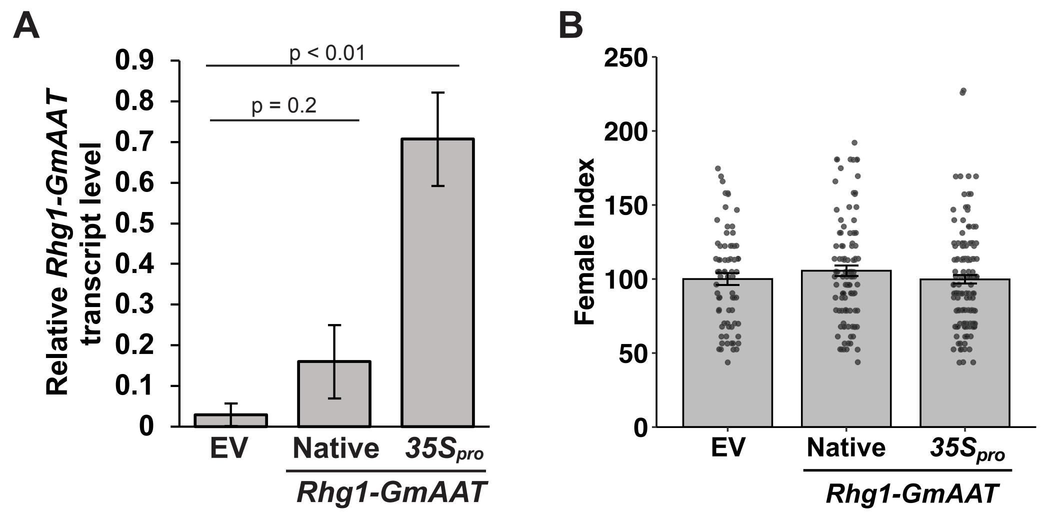


**Supplemental Figure S5. Overexpression of *Rhg1-GmAAT* in Arabidopsis does not enhance resistance against BCN.**
**(A)** Relative transcript abundance of *Rhg1-GmAAT* in stable transgenic Arabidopsis lines transformed with an empty vector (EV), a native *Rhg1-GmAAT* construct (native promoter, UTRs, exons, introns and terminator) or a *35S promoter:Rhg1-GmAAT CDS:Nos terminator* overexpression construct. Data were analyzed using one-tailed *t*-test and are presented as mean ± SE.
**(B)** Transgenic Arabidopsis lines described above were tested against BCN. Female index was calculated relative to EV. ANOVA did not show significant differences. Data are presented as mean ± SE.

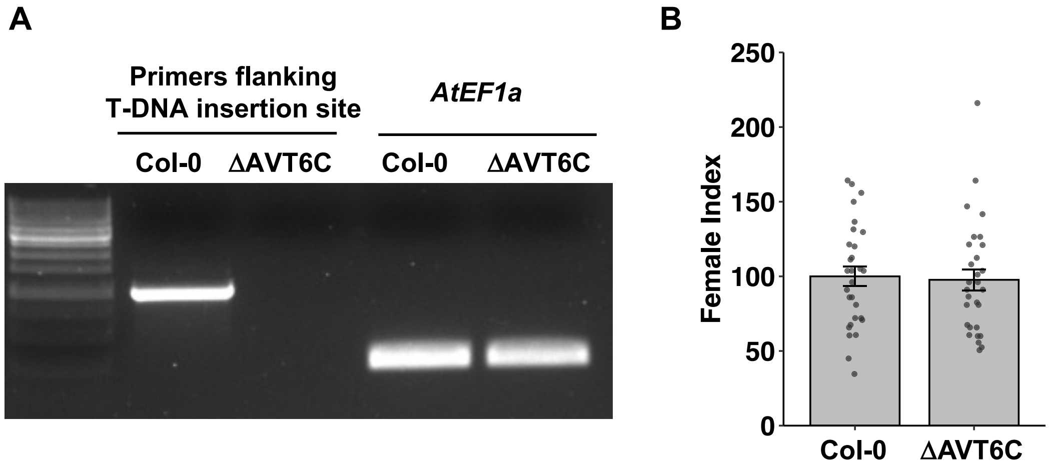


**Supplemental Figure S6. Knocking out *AVT6C* in Arabidopsis does not impact BCN resistance.**

**(A)** Verification of *AVT6C* knock-out in Arabidopsis line CS864997 (ΔAVT6C) obtained from Arabidopsis Biological Resource Center (ABRC). PCR was performed using primers flanking the T-DNA insertion site in the *AVT6C* gene, and genomic DNA extracted from Col-0 or ΔAVT6C plants as templates. A fragment of *AtEF1α* was amplified from the same genomic DNA templates as a control.

**(B)** The ΔAVT6C line was tested for resistance to BCN. Female index was calculated relative to Col-0. Data are presented as mean ± SE. Each bar contains four independent biological replicates. Each dot corresponds to data from an individual plant. A two-way ANOVA showed no significant differences between genotypes or replicates.

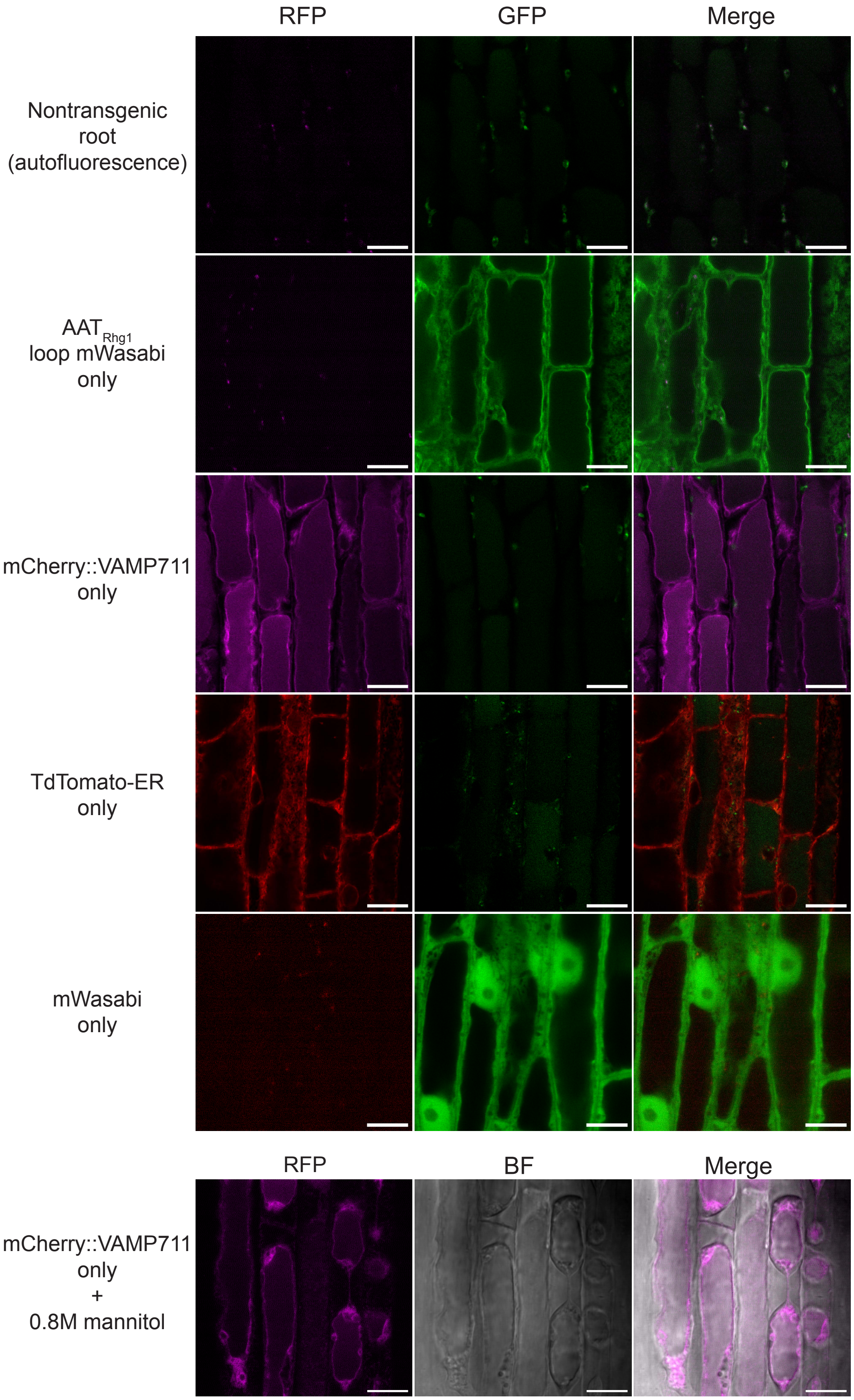


**Supplemental Figure S7. Confocal imaging controls confirm channel specificity.**

Representative confocal microscopy images of soybean roots showing, from top to bottom: non-transgenic roots (signal represents autofluorescence), $AAT_{Rhg1}$ loop mWasabi only, mCherry::VAMP711 only, TdTomato-ER only, soluble mWasabi only, and mCherry::VAMP711 only + 0.8M mannitol for plasmolysis. No signal bleed-through was detected between RFP and GFP channels. Scale bars = 10 μm.

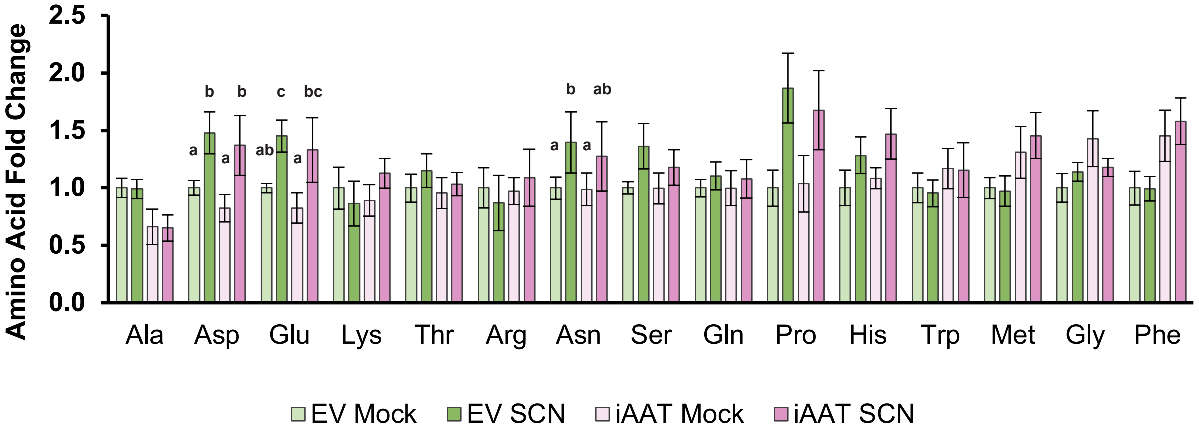


**Supplemental Figure S8. Amino acids unaffected by silencing *Rhg1-GmAAT* in the *rhg1-b* background.**
Quantification of amino acid levels in IL3025N (*rhg1-b*) empty vector (EV) or *Rhg1-GmAAT* RNAi-silenced (iAAT) transgenic soybean roots, either mock-treated or inoculated with SCN (HG 0) at 3 days post-inoculation (dpi). Amino acid levels were normalized as fold changes relative to EV-Mock. Data for each amino acid were analyzed using a linear model including group and replicate as fixed effects. Within each amino acid, bars sharing the same letter are not significantly different. Amino acids without letter annotations showed no significant differences. Data are presented as mean ± SE (*n* = 6, except Ala *n* = 3).

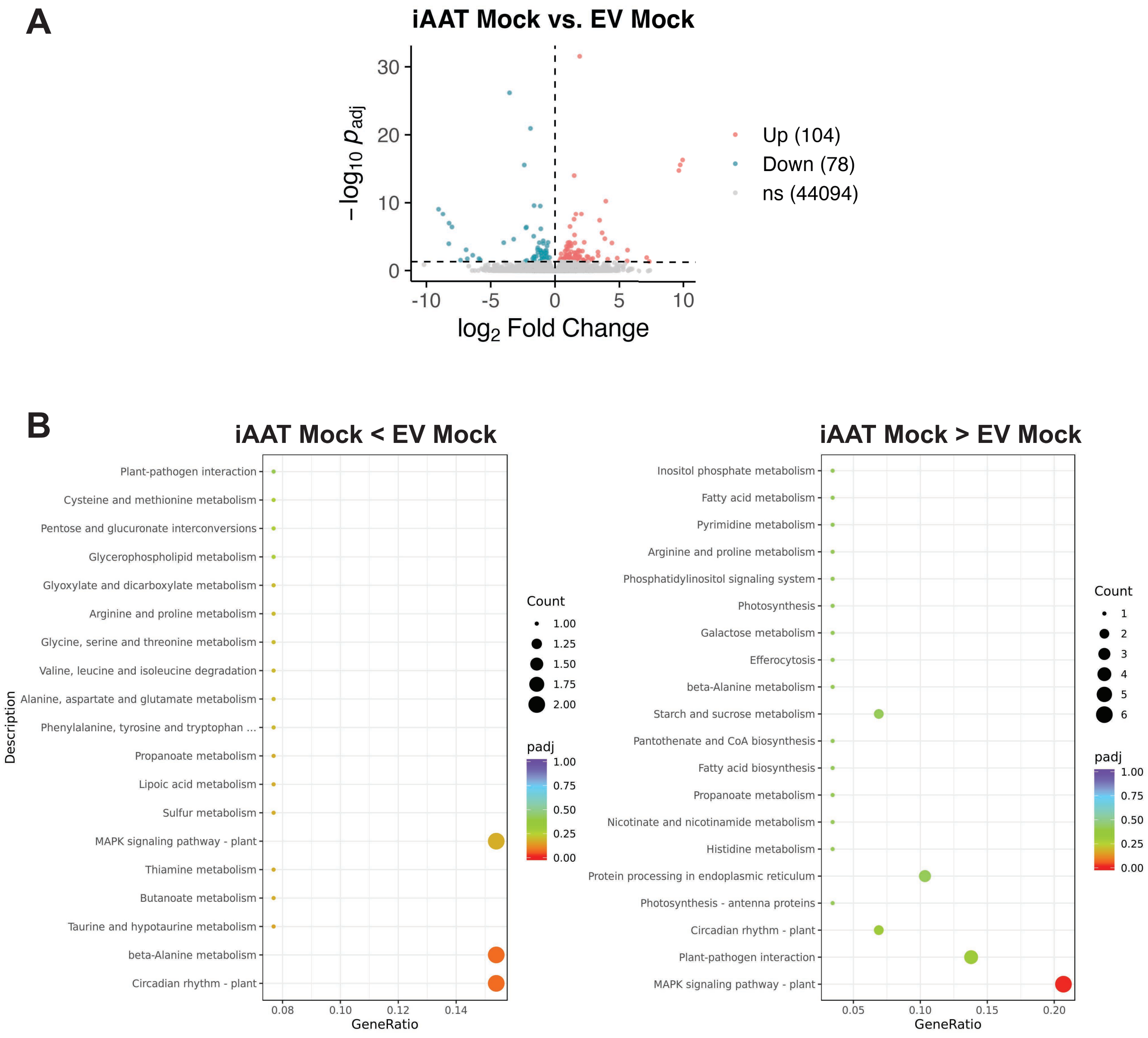


**Supplemental Figure S9. Differential gene expression between iAAT and EV under mock conditions.**

**(A)** Volcano plot showing differentially expressed genes (DEGs) between the mock-inoculated EV and iAAT roots. Each point represents a single gene. Genes with an adjusted *p*-value ($p_{adj}$) < 0.05 are highlighted.

**(B)** KEGG pathway enrichment analysis of DEGs identified in (A). Dot plots show the top 20 enriched KEGG pathways.

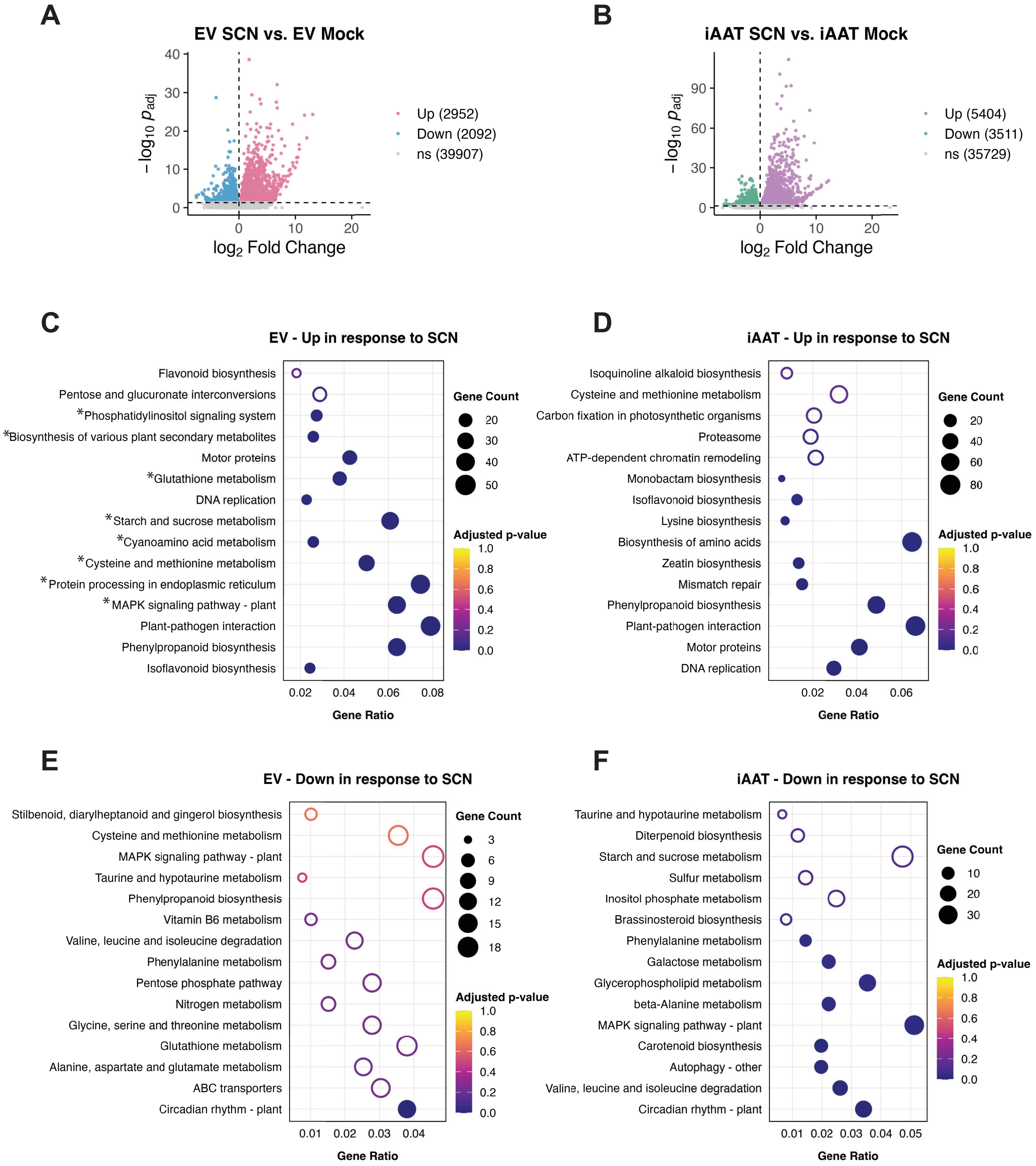


**Supplemental Figure S10. Differential gene expression between iAAT and EV in response to SCN.**

**(A, B)** Volcano plot showing differentially expressed genes (DEGs) between mock and SCN-inoculated conditions at 3dpi in the EV (A), or iAAT (B) roots. Each point represents a single gene. Genes with an adjusted *p*-value ($p_{adj}$) < 0.05 are highlighted. **(C-F)** KEGG pathway enrichment analysis of DEGs identified in (A) and (B). Dot plots show the top 15 enriched KEGG pathways based on upregulated genes in EV (C), upregulated genes in iAAT (D), downregulated genes in EV (E), and downregulated genes in iAAT (F) in response to SCN. Dot size reflects the number of DEGs associated with each pathway, and dot color represents enrichment significance. Filled circles indicate significantly enriched pathways ($p_{adj} < 0.05$), while empty circles represent non-significant enrichment. KEGG groups significantly upregulated by SCN in *rhg1-b* EV roots but not in iAAT roots are marked by asterisks in (C).

| Abundance (FPKM) | | | | | Abundance Difference ($p_{adj}$) | | | | |
|---|---|---|---|---|---|---|---|---|---|
| EV | | iAAT | | | SCN vs. mock | | iAAT vs. EV | | DEG in over-represented MAPK signaling KEGG |
| mock | SCN | mock | SCN | Gene | EV | iAAT | mock | SCN | |
| 11.1 | 10.4 | 11.7 | 9.2 | *GLYMA_03G245000* | 0.6451 | 0.0486 | 0.9999 | 0.7665 | 5'-3' exoribonuclease 4 |
| 15.9 | 16.3 | 17.6 | 13.3 | *GLYMA_19G242400* | 0.8839 | 0.033 | 0.9999 | 0.4143 | 5'-3' exoribonuclease 4 |
| 30.0 | 24.3 | 30.4 | 22.6 | *GLYMA_04G017500* | 0.1759 | 0.0264 | 0.9999 | 0.9128 | catalase |
| 64.4 | 44.7 | 67.7 | 41.5 | *GLYMA_14G223500* | 0.0509 | 0 | 0.9999 | 0.9106 | catalase |
| 40.1 | 31.9 | 45.2 | 28.5 | *GLYMA_06G017900* | 0.4203 | 0.0251 | 0.9999 | 0.8712 | catalase |
| 5.3 | 3.8 | 4.7 | 3.5 | *GLYMA_03G086000* | 0.2519 | 0.0263 | 0.9999 | 0.915 | copper-transporting ATPase HMA4 |
| 16.4 | 21.6 | 24.5 | 14.8 | *GLYMA_13G166200* | 0.0908 | 0.0225 | 0.4631 | 0.2779 | EIN3-binding F-box protein 1 |
| 40.4 | 30.9 | 48.2 | 25.8 | *GLYMA_04G066900* | 0.5174 | 0.0144 | 0.9999 | 0.8745 | EIN3-binding F-box protein 1 |
| 17.7 | 19.6 | 23.6 | 12.4 | *GLYMA_17G113900* | 0.8116 | 0 | 0.9999 | 0.1987 | EIN3-binding F-box protein 1 |
| 39.4 | 33.0 | 68.8 | 32.6 | *GLYMA_17G211000* | 0.4888 | 0.0001 | 0.1935 | 0.9888 | EIN3-binding F-box protein 1 |
| 27.6 | 23.6 | 43.3 | 27.1 | *GLYMA_14G116800* | 0.6602 | 0.0191 | 0.6401 | 0.8544 | EIN3-binding F-box protein 1 |
| 91.9 | 75.3 | 94.5 | 75.8 | *GLYMA_02G274600* | 0.1536 | 0.0036 | 0.9999 | 0.9962 | ETHYLENE INSENSITIVE 3-like 1 protein |
| 17.4 | 12.7 | 16.4 | 10.3 | *GLYMA_13G342500* | 0.2112 | 0.0031 | 0.9999 | 0.6183 | ETHYLENE INSENSITIVE 3-like 3 protein |
| 15.8 | 10.7 | 14.9 | 10.1 | *GLYMA_15G031800* | 0.0592 | 0.0002 | 0.9999 | 0.9363 | ETHYLENE INSENSITIVE 3-like 3 protein |
| 6.2 | 9.3 | 18.0 | 9.5 | *GLYMA_20G202200* | 0.2637 | 0.0238 | 0.0354 | 0.9888 | ethylene receptor 2 |
| 33.1 | 25.2 | 31.8 | 22.0 | *GLYMA_13G145100* | 0.092 | 0.0386 | 0.9999 | 0.8113 | ethylene-insensitive protein 2 |
| 1.6 | 1.9 | 12.1 | 3.1 | *GLYMA_13G123100* | 0.8095 | 0.0123 | 0.0168 | 0.6594 | ethylene-responsive transcription factor 1-like protein |
| 11.0 | 9.2 | 12.3 | 9.0 | *GLYMA_10G167300* | 0.4309 | 0.0407 | 0.9999 | 0.9795 | GAF/HisKA/CheY-like REC domain-containing protein,EIN4 |
| 17.1 | 13.3 | 16.8 | 13.4 | *GLYMA_08G115200* | 0.0735 | 0.0278 | 0.9999 | 0.9915 | mitogen-activated protein kinase 7 |
| 13.5 | 11.8 | 12.0 | 9.7 | *GLYMA_20G161500* | 0.1335 | 0.0247 | 0.9999 | 0.2158 | mitogen-activated protein kinase kinase kinase YODA |
| 50.7 | 37.6 | 65.9 | 37.4 | *GLYMA_13G076700* | 0.1631 | 0 | 0.8172 | 0.9947 | protein ETHYLENE INSENSITIVE 3 |
| 77.4 | 52.9 | 98.8 | 59.7 | *GLYMA_20G051500* | 0.0702 | 0 | 0.9709 | 0.8265 | protein ETHYLENE INSENSITIVE 3 |
| 40.4 | 41.5 | 45.8 | 28.7 | *GLYMA_04G205400* | 0.9698 | 0.0306 | 0.9999 | 0.2555 | protein kinase SRK2A |
| 34.2 | 43.9 | 50.1 | 34.9 | *GLYMA_08G005100* | 0.5652 | 0.0132 | 0.4379 | 0.8083 | protein kinase 3 SRK2A |
| 101.9 | 91.9 | 105.1 | 65.1 | *GLYMA_14G066400* | 0.7687 | 0.0199 | 0.9999 | 0.5167 | protein phosphatase 2C 37 |
| 57.8 | 50.4 | 68.5 | 43.5 | *GLYMA_17G052500* | 0.5465 | 0.0001 | 0.9999 | 0.7555 | protein phosphatase 2C 56 |
| 57.6 | 47.0 | 64.3 | 39.2 | *GLYMA_13G106800* | 0.4991 | 0.0063 | 0.9999 | 0.777 | protein phosphatase 2C 77 |
| 41.8 | 35.0 | 64.1 | 30.8 | *GLYMA_06G068400* | 0.5953 | 0.0014 | 0.4701 | 0.9041 | putative EIN3-binding F-box protein |
| 10.1 | 8.5 | 11.2 | 7.7 | *GLYMA_10G066000* | 0.6399 | 0.0461 | 0.9999 | 0.8931 | serine/threonine-protein kinase CTR1 |
| 9.3 | 8.4 | 8.5 | 6.2 | *GLYMA_07G178600* | 0.6985 | 0.0477 | 0.9999 | 0.454 | serine/threonine-protein kinase SRK2E |
| 29.2 | 22.2 | 30.4 | 23.6 | *GLYMA_11G058800* | 0.1204 | 0.0191 | 0.9999 | 0.927 | serine/threonine-protein kinase SRK2I |
| 16.9 | 12.6 | 14.5 | 10.5 | *GLYMA_11G072900* | 0.0903 | 0.0378 | 0.9999 | 0.7147 | uncharacterized LOC100527909 |

| | | | | | | | | | Additional DEG in "Unique Down in iAAT" |
|---|---|---|---|---|---|---|---|---|---|
| 81.4 | 33.7 | 60.0 | 20.6 | *GLYMA_07G264200* | 0.074 | 0 | 0.9999 | 0.6406 | 1-aminocyclopropane-1-carboxylate oxidase |
| 11.5 | 0.7 | 14.0 | 2.5 | *GLYMA_08G169100* | 0.0743 | 0.0486 | 0.9999 | 0.6078 | 1-aminocyclopropane-1-carboxylate oxidase homolog 11-like |
| 37.3 | 18.7 | 44.4 | 23.3 | *GLYMA_08G351866* | 0.3567 | 0.0461 | 0.9999 | 0.8431 | 1-aminocyclopropane-1-carboxylate oxidase homolog 12 |
| 34.6 | 27.5 | 41.1 | 20.0 | *GLYMA_07G168500* | 0.4335 | 0.004 | 0.9999 | 0.4832 | 1-aminocyclopropane-1-carboxylate oxidase homolog 4 |
| 10.4 | 7.6 | 13.1 | 7.6 | *GLYMA_07G021000* | 0.2581 | 0.0233 | 0.9999 | 0.9978 | AP2-like ethylene-responsive transcription factor At1g16060 |
| 37.4 | 25.6 | 43.2 | 21.5 | *GLYMA_06G042100* | 0.4663 | 0.0116 | 0.9999 | 0.8494 | dehydration-responsive element binding protein 2 |
| 1.6 | 1.3 | 1.5 | 0.6 | *GLYMA_09G052900* | 0.8939 | 0.035 | 0.9999 | 0.7194 | DNA-binding domain-containing protein |
| 26.6 | 22.8 | 27.6 | 17.5 | *GLYMA_10G058300* | 0.4531 | 0.0004 | 0.9999 | 0.515 | ethylene-insensitive protein 2 |
| 10.3 | 6.9 | 12.6 | 6.0 | *GLYMA_11G167200* | 0.4593 | 0.0001 | 0.9999 | 0.9124 | ethylene-overproduction protein 1 |
| 5.2 | 3.0 | 4.7 | 2.9 | *GLYMA_18G059700* | 0.136 | 0.0002 | 0.9999 | 0.9428 | ethylene-overproduction protein 1 |
| 78.6 | 64.2 | 144.0 | 80.8 | *GLYMA_13G236500* | 0.4711 | 0.0219 | 0.3586 | 0.7022 | ethylene-responsive element binding factor 4 |
| 15.0 | 7.8 | 14.8 | 9.5 | *GLYMA_08G137600* | 0.0652 | 0.0165 | 0.9999 | 0.7601 | ethylene-responsive transcription factor |
| 42.6 | 43.3 | 73.0 | 36.3 | *GLYMA_13G274100* | 0.9752 | 0 | 0.5714 | 0.7325 | ethylene-responsive transcription factor 4 |
| 14.6 | 10.9 | 15.9 | 10.3 | *GLYMA_11G239200* | 0.241 | 0.0078 | 0.9999 | 0.9585 | ethylene-responsive transcription factor ERF060-like |
| 0.1 | 1.1 | 6.5 | 1.4 | *GLYMA_10G036200* | 0.0315 | 0.0307 | 0.001 | 0.9245 | ethylene-responsive transcription factor ERF096 |
| 2.4 | 3.7 | 10.7 | 2.2 | *GLYMA_13G122500* | 0.5031 | 0.001 | 0.033 | 0.7523 | ethylene-responsive transcription factor ERF096 |
| 4.3 | 3.0 | 3.4 | 0.9 | *GLYMA_10G223200* | 0.7097 | 0.0022 | 0.9999 | 0.1836 | ethylene-responsive transcription factor ERF110 |
| 1.1 | 0.2 | 2.9 | 0.5 | *GLYMA_13G060600* | 0.0552 | 0.0308 | 0.76 | 0.7739 | ethylene-responsive transcription factor ERN2 |
| 15.7 | 10.3 | 21.6 | 7.3 | *GLYMA_04G041200* | 0.3871 | 0 | 0.9999 | 0.683 | ethylene-responsive transcription factor RAP2-1 |
| 23.0 | 18.3 | 25.4 | 10.0 | *GLYMA_12G073300* | 0.675 | 0.0028 | 0.9999 | 0.2329 | ethylene-responsive transcription factor RAP2-7 |
| 7.3 | 5.7 | 6.9 | 4.7 | *GLYMA_09G177600* | 0.2757 | 0.0405 | 0.9999 | 0.7396 | probable serine/threonine-protein kinase SIS8 |
| 13.6 | 12.8 | 21.6 | 7.8 | *GLYMA_14G182700* | 0.8894 | 0.0005 | 0.911 | 0.3217 | serine/threonine-protein kinase CTR1 |
| 14.1 | 11.8 | 14.2 | 11.0 | *GLYMA_20G229500* | 0.1194 | 0.0191 | 0.9999 | 0.8317 | serine/threonine-protein kinase EDR1 |
| 15.0 | 14.7 | 25.2 | 15.7 | *GLYMA_02G066200* | 0.967 | 0.0364 | 0.8102 | 0.9456 | uncharacterized LOC100796480 |

FPKM: 0 – 72 – 144

$p_{adj}$: 0.001 – 0.3 – 1.0

**Supplemental Figure S11. Genes uniquely downregulated in iAAT in response to SCN that are associated with MAPK signaling and ethylene responses.**
Top: Transcripts significantly downregulated at SCN infection sites in *Rhg1-GmAAT*-silenced roots (iAAT) but not in SCN-resistant *rhg1-b* (EV) roots, for over-represented KEGG group "MAPK Signaling Pathways – Plants". Bottom: Additional genes uniquely downregulated in iAAT but not in EV roots in response to SCN, that carry ethylene-associated annotation but are not placed by KEGG into the above group.

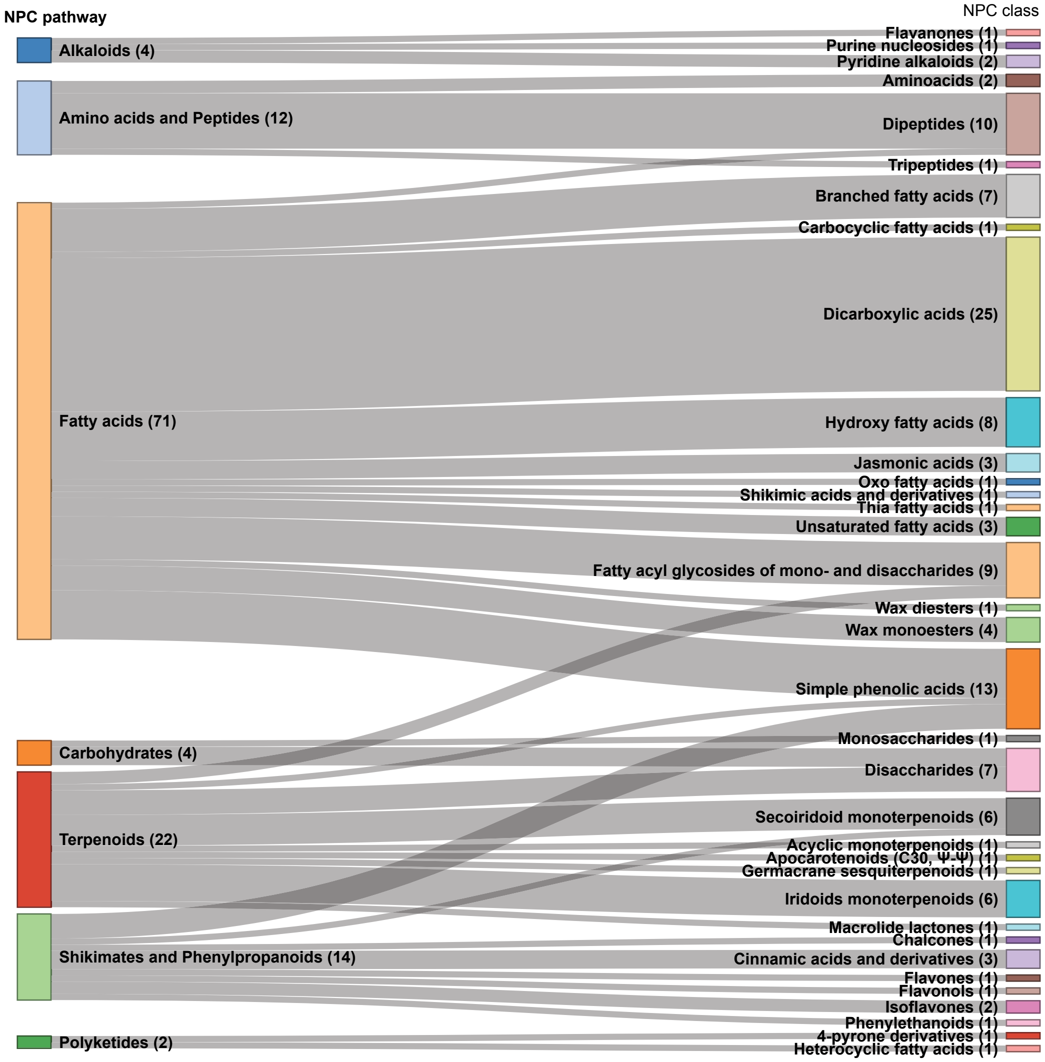


**Supplemental Figure S12. Composition of differentially abundant metabolites detected in RP-negative mode.**

Differentially abundant metabolites (fold change > 2 or < 0.5; *t*-test $p < 0.05$, as defined in Fig. 7B and 7C) with available NPC annotations were grouped by NPC pathways and classes. Bars represent the distribution of these annotated metabolites.
